# Histone deacetylase inhibitors butyrate and bufexamac inhibit *de novo* HIV-1 infection in CD4 T-cells

**DOI:** 10.1101/2020.04.29.067884

**Authors:** Lin Chen, Ariane Zutz, Julia Phillippou-Massier, Tim Liebner, Oliver T. Keppler, Chunaram Choudhary, Helmut Blum, Christian Schölz

## Abstract

While current combined antiretroviral therapy (cART) allows control of HIV replication in patients and effectively suppresses plasma viral loads, it is unable to target latent reservoirs, which are responsible for virus rebound after discontinuation of therapy. Several histone deacetylase inhibitors (HDACIs) have been shown to target reservoirs and to reactivate latent HIV. While this effect is highly desired, it carries the risk that HIV-1 may be reactivated in tissue compartments were cART concentrations are insufficient and thus leading to *de novo* infections in this sites. To address this concern, we evaluated the effect of different HDACIs for their ability to reverse HIV latency and to modulate *de novo* infections. Two of the inhibitors, sodium butyrate and bufexamac, significantly inhibited *de novo* HIV-1 infection in activated CD4^+^ T-cells. Transcriptome and proteome analysis indicated global changes of protein abundancies, exhibited reduced proliferation of CD4^+^ T-cells, and revealed butyrate-based proteasomal degradation of EP300, an important factor for HIV-1 replication. Our results disclose new potential treatment strategies and minimizes the concern of potential reservoir reseeding by HDACIs.

## Introduction

Combined antiretroviral therapy (cART) allows control of HIV-1 replication in infected individuals, suppresses viral loads to undetectable levels, and reconstitutes immune function, which effectively reduces morbidity and mortality. However, during the acute phase of infection, HIV-1 establishes also transcriptionally silent, long-lived reservoirs of replication-competent provirus, allowing viral rebound upon cessation of antiviral treatment [1,2].

These reservoirs stay largely unaffected by cART and are the major obstacle for virus eradication. Thus, one of the major goals in HIV therapy is the elimination of latent reservoirs.

Despite its clinical relevance, the molecular mechanisms that are responsible for the establishment and maintenance of HIV latency remain largely undefined. Several studies have shown that proviral integration in epigenetically silenced heterochromatin regions in combination with other factors, like regulators of transcription, might lead to latency in long-lived memory cells [3–5]. More detailed analyses disclosed a strong dependency on histone deacetylases (HDACs) for sustained HIV-1 latency. HDACs remove acetyl-groups from acetylated lysines and thus are directly involved in the deacetylation of histone sites as well as the corresponding condensation of the chromatin, which inhibits transcription of viral genes [6,7]. Moreover, transcriptional repressors, as well as several (co-)repressors and transcription factors (TFs) actively recruite HDACs to the retroviral promoter sites, known as long terminal repeats (LTR) [8–14]. Overall, 44 % of all known human deacetylases including sirtuins have been shown to play pivotal roles during the HIV life cycle supported also by around 140 acetylation-related interactions (see also [15]).

Consistent with the importance of HDACs for establishment and maintenance of latency it is not surprising that histone deacetylase inhibitors (HDACIs) are able to interfere with the mechanisms for latency perpetuation as shown in both ex vivo and in vivo studies [16–20]. Within recent years, several approaches have been undertaken to further characterize and destroy latent reservoirs. One of these methods, the so called “kick and kill” strategy aims at the specific reactivation of the virus by chemical compounds, in particular HDACIs, with subsequent elimination by cART [19].

A clinical concern is that reactivation of virus might lead to *de novo* infections in tissues where cART concentrations are suboptimal and thus a reseeding of the latent reservoir. This hypothesis is also supported by a study demonstrating that high concentrations of romidepsin, an HDACI, which very effectively reactivates latent HIV-1, support infection of CD4^+^ T-cells with HIV-1 [21]. Moreover, so far a significant decrease of the size of the latent HIV-1 reservoir in patients by kick and kill approaches has not been observed [22] and might be related to reseeded reservoirs. So far only very limited information is available about possible side effects of HDACIs as well as the effect of HDACIs on *de novo* HIV infections [23–25].

To comprehensively determine the possibility of HDACI-induced enhancement of *de novo* HIV-1 infections in relevant target cells, we tested a broad panel of HDACIs for (i) their ability to reverse HIV latency and for (ii) their impact on viral infection in human primary activated CD4^+^ T-cells. Using this *ex vivo* experimental system, we further identified two novel compounds, sodium butyrate and bufexamac, which profoundly restricted *de novo* HIV infections. Detailed analysis of the mode of action of both drugs revealed global intracellular changes on transcriptome and proteome level. In addition, we identified the histone acetyltransferase EP300, which has been shown to be important for the transcriptional activity of the viral protein Tat [26,27], as a substantial intracellular target of butyrate treatment, leading to its proteasomal degradation.

Our results might aid the development of novel potential treatment strategies against HIV-1 and minimize the worry of reservoir reseeding by HDACIs in general.

## Results and Discussion

### Reactivation of latent HIV-1 by HDACIs

To comprehensively study the effect of HDACIs on *de novo* HIV-1 infections, we selected 15 HDACIs, covering the entire selectivity range for human deacetylases including sirtuins, although with varying specificity profiles (**Figure EV1**). First, we evaluated the capability of the different HDACIs for reversal of HIV-1 latency. Therefore, we challenged J-Lat-cells, clones 8.4 and 10.6, derived from Jurkat T-cells that had been transduced with near full-length, *env*- (non-functional due to frameshift) and *nef*-defective HIV R7, that carries a *gfp*-reporter cassette in the *nef* locus [28], with increasing concentrations of the respective compounds. GFP expression indicates LTR-driven early HIV-1 gene expression. Solvent only treatment (DMSO or H_2_O) of J-Lat-cells was used as a control and typically yielded on average 6-7 % GFP-positive cells, indicating this cell model’s background. 24 h post treatment, cells were analyzed for GFP expression by flow cytometry (**Figures 1A, EV2**). Reactivation of HIV by selected compounds was further monitored by fluorescence microscopy, as illustrated in **Figure 1B**. As expected for HDACIs, most of the compounds were able to reactivate HIV-1 efficiently and allowed us to cluster them into three groups (**Figures 1A, EV2**). The first group comprised inhibitors such as romidepsin, apicidin, TSA, and LBH589 (panobinostat), which are known to target class I and II HDACs, and which displayed efficient reactivation of HIV within a low concentration range (10^−5^ to 10^−7^ M). The second group triggered reactivation at significantly higher concentrations, i.e. in the mM range, as shown for e.g. valproate or sodium butyrate (Sb). For the last group no significant HIV-1 GFP reactivation signal was obtained at the concentrations tested. Inhibitors of this group included sirtuin inhibitors splitomicin and nicotinamide and HDAC6 inhibitors bufexamac (Bu) and tubacin, which are known to have less effect on overall histone acetylation [29]. Thus, our data strongly indicate that in particular class I and II HDACs are modulating HIV-1 gene expression and latency and support previous studies for individual HDACIs.

**Figure 1:**
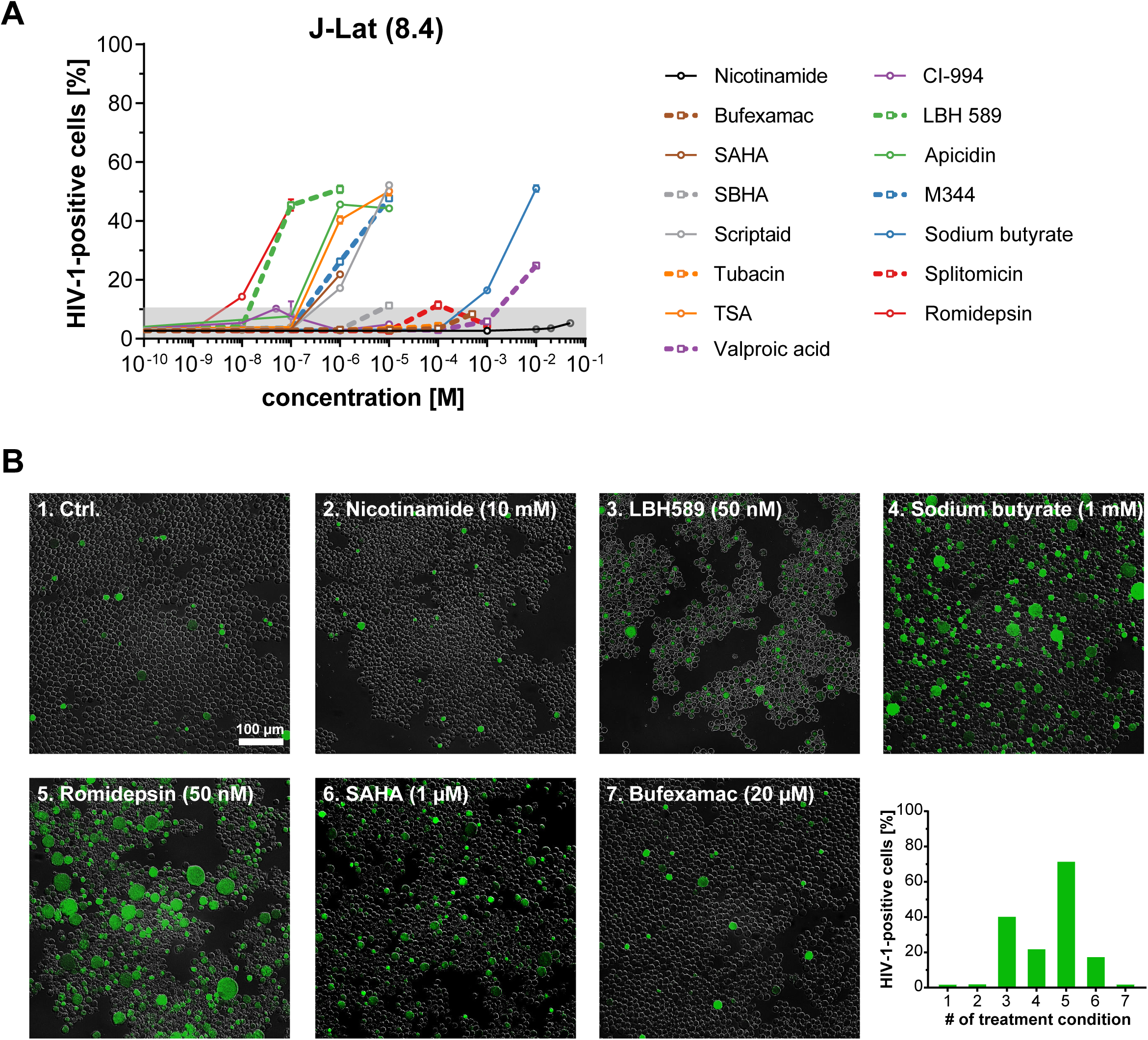
Reactivation of latent HIV by HDAC inhibitors. A) J-Lat cells (clone 8.4) were treated with increasing inhibitor concentrations for 24 h. Subsequently, percentage of reactivation was determined by GFP-based flow cytometry. Data are derived from three independent experiments. Error bars show SD. See also Figure S2. B) Fluorescence microscopy-based monitoring of reactivation-accompanied expression of GFP. Shown are representative images of untreated control cells (1.), and six differently treated examples (2.-7.). Lower right graph displays the corresponding percentages of GFP positive cells for the shown samples 1.-7. as measured by flow cytometry.

### Effect of HDACI treatment on HIV-1 *de novo* infection

After verifying the efficacy of the HDACIs in this study for latency removal, we addressed the concern that latency reversing agents (LRAs) might also function as replication enhancers, which would promote *de novo* infection with HIV-1 and thus a reseeding of the latent reservoir. To resemble closely the clinical conditions, we made use of human primary, activated CD4^+^ T-cells, isolated from peripheral blood of healthy donors, and full-length, replication-competent HIV-1_NL4-3_ strain. In contrast to several other studies, this strain has not been pseudotyped with Vesicular stomatitis virus glycoprotein (VSV-G) and thus mirrors more closely the *in vivo* situation. We first determined the toxicity of the HDACIs in activated CD4^+^ T-cells (purity and activation status was controlled by flow cytometry; **Figure EV3**) following a treatment period of 72 h to simulate the duration of the subsequent infection experiments (**Figure EV4**). Employed dose ranges covered physiological and scientifically used concentrations (e.g. ∼4 mM butyrate [30], 0.5 µM SAHA [31], 30 nM LBH589 [32]). A toxicity of 20 % (lethal dose 20 %; LC_20_) was tolerated for each compound and used as maximal concentration for all infection experiments (**Table 2**). Notably and as shown for the treatment with romidepsin, cells from different blood donors displayed an expected level of variability, which could be mitigated by using 4-donor pools in subsequent experiments (**Figure EV4**). Next, we explored whether the LC_20_ concentrations are still in a functional range to efficiently affect enzymatic activity and to trigger upregulation of cellular acetylation. Using both, a PAN-acetyl staining-based flow cytometry readout (**Figure EV5A**) and quantification of histone H3 acetylation by ELISA (**Figure EV5B**), we could confirm the efficacy of the used drugs and concentrations. While all inhibitors caused a global increase in acetylation, albeit at varying intensities, histone H3 acetylation was mainly elevated by class I and IIa compounds. HDACIs targeting either class IIb or sirtuins, showed expectedly, no or very low effects on histone acetylation (**Figure EV5B**). Finally, we explored the effect of HDACI treatment on *de novo* infection of 4-donor pools of activated CD4^+^ T-cells. Cells were pretreated with HDACIs for 24 h and subsequently infected with full-length, replication-competent HIV-1_NL4-3_. After cultivation for 48 h, the number of infected cells was analyzed by intracellular p24 antigen staining and flow cytometry (**Figures 2A-C**). Interestingly and despite of the previous latency removal experiments, none of the tested compounds was able to enhance *de novo* infections in activated CD4^+^ T-cells (**Figure 2C**). In general, HDACI treatment did not or only moderately affect *de novo* infection levels in most cases. However, treatment with two of the inhibitors, Sb and Bu, both shown to target very different HDACs, i.e. class I [33] HDACs and HDAC6/10 [34], respectively, caused an unexpected, highly significant reduction of HIV infection levels, leading to a decrease of p24-positive cells of around 70-75% compared to vehicle-treated control cells (**Figure 2C**). Similar findings were made for the R5-tropic HIV-1 transmitted founder (T/F) strain CH058 [35], demonstrating that the observed reduction was independent of the coreceptor usage and was observed for a molecular clone that closely resembles primary isolates (**Figure 2D**). Findings were compound-specific and did not correlate with the compounds’ target specificity (e.g. tubacin, another HDAC6-specific inhibitor and LBH589, another class I/II inhibitor, respectively, did not negatively affect intracellular p24 levels in our screen). Overall, these data disprove the potential risk of enhanced *de novo* infections at least in CD4^+^ T-cells and highlight the two HDACIs Sb and Bu as potential novel anti-HIV drugs.

**Table 1:** Key resources table.

| Reagent type (species) or resource | Designation / Name | Source or reference | Identifiers | Additional information |
| --- | --- | --- | --- | --- |
| Bacteria strain | Stbl II | Thermo Fisher Scientific | #10268019 |  |
| Cell line | Hek293T | DSMZ | ACC-635 |  |
| Cell line | TZM-bl | NIH AIDS Reagent Programme | 8129 |  |
| Cell line | J-Lat (clone 10.6) | NIH AIDS Reagent Programme | 9849 | HIV-1 latent T-cell clone |
| Cell line | J-Lat (clone 8.4) | NIH AIDS Reagent Programme | 9847 | HIV-1 latent T-cell clone |
| Primary cells | CD4<sup>+</sup> T-cells | Isolated from healthy donors |  | Confirmed by ethics committee of the LMU; Germany |
| Antibody cocktail | RosetteSep™ Human CD4+ T-cell Enrichment Cocktail | Stemcell Technologies | #15022 | Isolation of CD4<sup>+</sup> T-cells |
| Recombinant DNA reagent | pNL4-3 | NIH AIDS Reagent Program | #114 | HIV-1 expresssion vector |
| Recombinant DNA reagent | pCH058.c/2960 | NIH AIDS Reagent Program | #11856 | HIV-1 Transmitted/Founder expression vector |
| Recombinant DNA reagent | pMM310 | NIH AIDS Reagent Program | #11444 | β-lactamase-Vpr expression vector |
| FACS Antibody | KC57-FITC | Beckman Coulter | #6604665 | Flow cytometry |
| FACS Antibody | BD Tritest™ CD4/CD8/CD3 | BD Biosciences | #340298 | Flow cytometry |
| FACS Antibody | Anti-Human CD4 FITC/CD69 PE/CD3 PerCP | BD Biosciences | #340365 | Flow cytometry |
| FACS Antibody | Anti-Human-CD25-APC | BD Biosciences | #555434 | Flow cytometry |
| FACS Antibody | Anti-Human-CD184 (CXCR4)-APC | BD Biosciences | #555976 | Flow cytometry |
| FACS Antibody | Anti-Human-CD195 (CCR5)-FITC | BD Biosciences | #555992 | Flow cytometry |
| FACS Antibody | PE anti-Acetylated Lysine Antibody | Biolegend | # 623404 | Global acetylation test |
| FACS Antibody | PE Mouse IgG2b, $\kappa$ Isotype Ctrl Antibody | Biolegend | # 400314 | Global acetylation test; isotype control |
| Buffer | Perm/Wash buffer | BD Biosciences | #554723 | Flow cytometry staining buffer |
| Viability Test | CellTiter-Glo® LuminescentT-cell Viability Assay | Promega | G7570 | Viability testing |
| Acetyl ELISA | PathScan® Acetylated Histone H3 Sandwich ELISA | Cell Signaling Technology | #7232C | Histone H3 acetylation |
| RT Activity Kit | Reverse Transcriptase Assay, colorimetric | Sigma Aldrich | #11468120910 | RT activity |
| C18 matrix | SEP-Pak Classic | Waters | WAT051910 | Peptide purification |
| Isobaric labels | TMTduplex™ Isobaric Mass Tagging Kit | Thermo Fisher Scientific | #90063 | TMT labeling of peptides |
| Fractionation Kit | High pH Reversed-Phase Peptide Fractionation Kit | Pierce (Thermo Fisher Scientific) | #84868 | Fractionation of peptides |
| Co-IP antibody | p300 Recombinant Mouse Monoclonal Antibody (2033) | Invitrogen | #730005 | Co-IP |
| Immunoblot antibody | p300 Monoclonal Antibody (NM-11) | Invitrogen | #337600 | Immunoblotting |
| Immunoblot antibody | Anti-vinculin antibody | abcam | #ab130007 | Immunoblotting |
| Magnetic beads | SureBeads Protein G Magnetic Beads | Bio-Rad | #161-4023 | Co-IP |
| Enzyme | SensoLyte® | AnaSpec | #AS-72172 | EP300 activity test |
| reactivity test | HAT (p300)<br>Assay Kit<br>`Fluorimetric` |  |  |  |

**Table 2:** Histone deacetylase inhibitors used in this study and determined LC_20_ values.

| HDACI | CAS number | Chemical name | Alternative name | Solvent | LC<sub>20</sub> |
| --- | --- | --- | --- | --- | --- |
| Apicidin | 183506-66-3 | Cyclo[(2S)-2-amino-8-oxodecanoyl-1-methoxy-L-tryptophyl-L-isoleucyl-(2R)-2-piperidinexcarbonyl] | OSI 2040 | DMSO | 0.09 µM |
| Bufexamac | 2438-72-4 | 2-(p-Butoxy-phenyl)aceto-hydroxamic acid |  | DMSO | 0.04 mM |
| CI-994 | 112522-64-2 | 4-(acetyl-amino)-N-(2-aminophenyl)-benzamid | Tacedinaline | DMSO | 3.33 µM |
| LBH589 | 404950-80-7 | (E)-N-hydroxy-3-(4-((2-(2-methyl-1H-indol-3-yl)ethylamino)methyl)phenyl)acrylamide | Panobinostat | DMSO | 0.01 µM |
| M344 | 251456-60-7 | 4-(Dimethylamino)-N-[7-(hydroxyamino)-7-oxoheptyl]-benzamide, MS 344, D237, N-Hydroxy-7-(4-dimethyl-aminobenzoyl)-aminoheptanamide | D237, MS 344 | DMSO | 0.75 µM |
| Nicotinamide | 98-92-0 | Pyridine-3-carboxylic acid amide | Niacinamide, Vitamin B3, Vitamin PP | H<sub>2</sub>O | 6.76 mM |
| Romidepsin | 128517-07-7 | (E)-(1S,4S,10S,21R)-7-[(Z)-ethylidene]-4,21-diisopropyl-2-oxa-12,13-dithia-5,8,20,23-tetraazabicyclo[8.7.6]tricos-16-ene-3,6,9,19, 22-pentone | Istodax, FK228, FR901228, Depsipeptide, NSC 630176 | DMSO | 1.29 nM |
| SAHA | 149647-78-9 | N'-hydroxy-N-phenyloctanediamide | Vorinostat, Zolinza | DMSO | 0.78 µM |
| SBHA | 38937-66-5 | N,N'-Dihydroxyoctanediamide | Suberic bis-Hydroxamic Acid | DMSO | 0.79 µM |
| Scriptaid | 287383-59-9 | 6-(1,3-Dioxo-1H, 3H-benzo[de]isoquinolin-2-yl)-hexanoic acid hydroxyamide | GCK 1026 | DMSO | 0.70 µM |
| Sodium butyrate | 156-54-7 | Butyric acid sodium salt | Butyric acid sodium salt | H<sub>2</sub>O | 1.95 mM |
| Splitomicin | 5690-03-9 | 1,2-Dihydro-3H-naphtho[2,1-b]pyran-3-one | 1-Naphthalenepropionic Acid | DMSO | 224.36 µM |
| TSA | 58880-19-6 | (2E,4E,6R)-7-[4-(dimethylamino)phenyl]-N-hydroxy-4,6- | Trichostatin A | DMSO | 0.24 µM |
|  |  | dimethyl-7-oxohepta-2,4-dienamide |  |  |  |
| Tubacin | 537049-40-4 | N-[4-[(2R,4R,6S)-4-[(4,5-diphenyl-1,3-oxazol-2-yl)sulfanylmethyl]-6-[4-(hydroxymethyl)phenyl]-1,3-dioxan-2-yl]phenyl]-N'-hydroxyoctane-diamide | | DMSO | 7.25 μM |
| Valproic acid | 99-66-1 | 2-propylpentanoic acid | Depakine, Depakene, Ergenyl, Dipropylacetic acid, Mylproin, Convulex, Valproic Acid, Myproic Acid | H<sub>2</sub>O | 1.34 mM |

**Figure 2:**
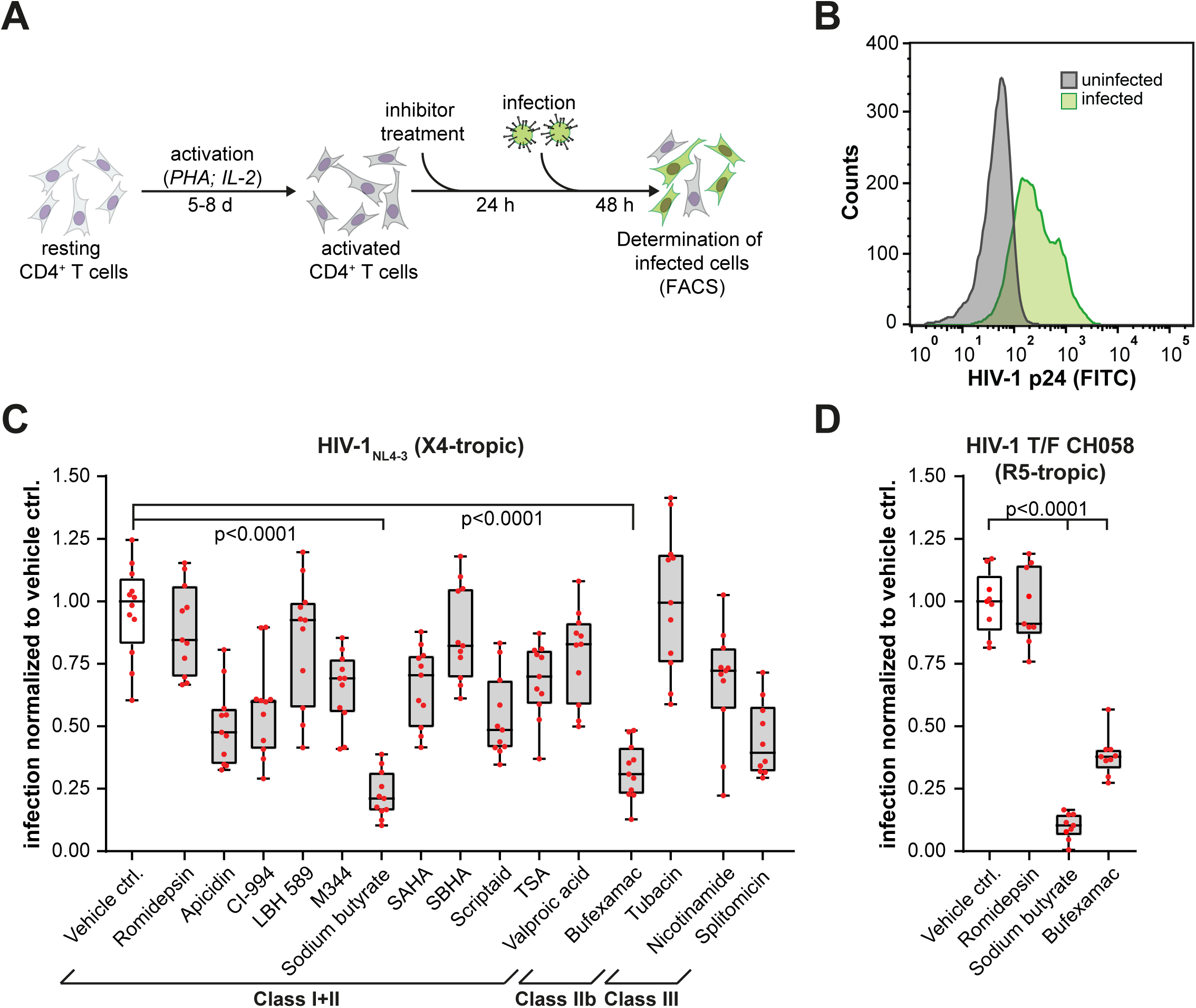
Effect of HDACIs on *de novo* HIV-1 infection. A) Experimental design. Isolated, primary CD4^+^ T-cells were activated by PHA and IL-2 treatment for 5-8 days. Derived activated T-cells were treated with HDACIs for 24 h and subsequently infected with HIV-1_NL4-3_. 48 h post infection, cells were stained for the viral p24 capsid protein and analyzed by flow cytometry. B) Representative FACS histogram plot showing the increase of p24 capsid protein in infected cells (green). Uninfected cells were used as control (grey) C) Box-Whisker plots representing the HIV-1_NL4-3_ infection levels in CD4^+^ T-cells in dependency of the respective HDACI treatment. All data were normalized to the median of vehicle control samples. Boxes represent the lower and upper quartiles, whiskers show the minimum and maximal values, and the line inside the box indicates the median. Individual data points are presented as red dots. Sample size: n ≥ 10. Given p-values show statistically difference (ANOVA-based Dunnetts’ multiple comparison test) between vehicle control and sodium butyrate and bufexamac treated cells. D) Graph similar to C except infection with R5-tropic HIV-1 T/F CH058 virus. Sample size: n = 9.

### Transcriptomic analysis of Sb- and Bu-treated cells

in order to better understand the underlying molecular mechanism of both HDACIs on a global level, we performed transcriptome and proteome analysis of Bu- and Sb-treated cells. For transcriptomic analysis, comprehensive mRNA profiles of samples 48 h post treatment (Sb n=3, Bu n=3, Ctrl. n=3) were generated by random-primed cDNA sequencing (RNA-Seq) with reduced representation of ribosomal RNA. The average depth of mapped reads was around 22 million reads. In total, 15.396 transcripts for Sb-treated (**Figure 3A**) and 12.833 transcripts for Bu-treated cells (**Figure 3B**) were identified (transcripts which failed statistical analysis (DESeq: no q-value) were excluded), of which >91 % could be annotated to ENSEMBL gene IDs. For sodium butyrate 3.585 differential expressed genes (DEGs) and for bufexamac 1.708 DEGs, respectively, were found (DESeq; FDR: 0.01) (see also **Supplemental data file**), of which 1.342 genes were significantly upregulated (log_2_ fold change (log_2_fc) ≥1) and 814 genes were downregulated (log_2_fc ≤-1) in Sb-treated samples and 61 genes were upregulated and 261 genes downregulated in Bu-treated samples, respectively (**Figures 3A, 3B, EV6A**). Remarkably, despite the different target specificity of both compounds, the differentially expressed genes showed good correlation with an R^2^-value of 0.76 (**Figure EV6A**). Moreover, comparison with the NCBI HIV Human Interaction Database (HHID) [36] showed that ∼25% (Sb) and ∼34% (Bu) of regulated genes, respectively, have been associated previously with HIV infection [36]. Evaluation of the 20 most up- and down-regulated transcripts (**Figure 6B**) displays increased abundancies for genes associated with signaling, such as TGFα and LRP6. Interestingly, downregulated genes comprise several cytokines/interleukins such as CXCL8 (IL8), which is known to promote HIV replication in human monocyte-derived macrophages [37] and IL4, which stimulates expression of HIV via a transcriptional activation mechanism [38] and additionally plays an important role in T-cell activation, proliferation and differentiation [39]. Gene ontology (GO) and pathway analysis using g:Profiler [40] identified gene sets related to “mitosis”, “G1/S-” and “G2/M-transition” or “DNA double strand break repair” being depleted in Sb- and Bu-challenged samples (**Figure 3C**), whereas genes associated with the cytoskeleton were exclusively enriched in cells challenged with Sb (**Figure EV6C**). Moreover, pathway analysis indicated significant deregulation of the KEGG pathways “cell cycle” and “Fanconia anemia pathways” by both inhibitors (**Figures 3C, EV7**). Data provide detailed insights into the mode of action of Sb and Bu, thus helping to understand their impact on HIV-1 infection. Moreover, results are also in line with earlier observations, showing that most HDACIs are able to block cellular proliferation.

**Figure 3:**
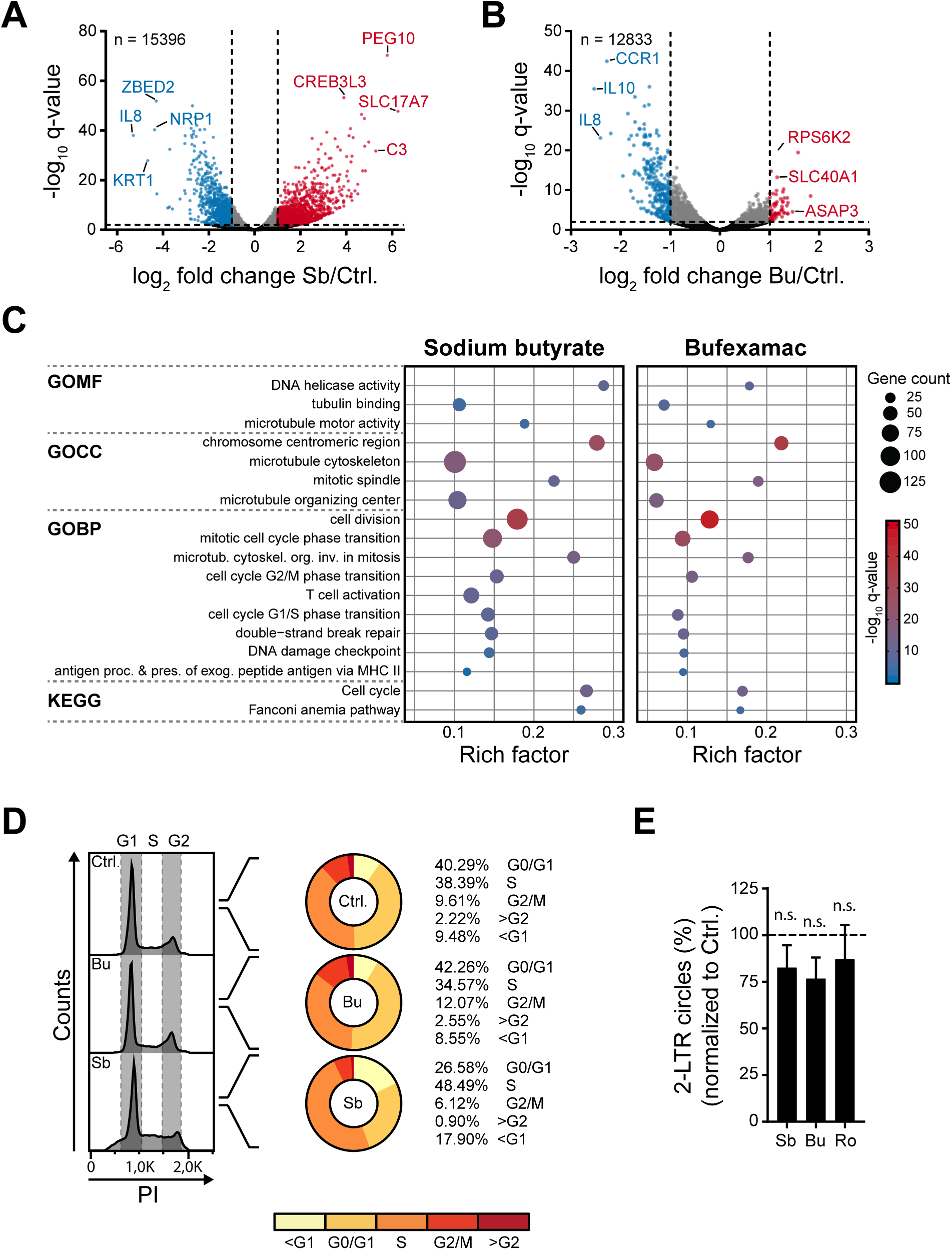
Transcriptome analysis of Sb- and Bu-treated cells. A and B) Volcano plots of identified transcripts showing the log_2_ fold change of abundance between inhibitor (A: Sb, B: Bu) and vehicle control and the associated q-value for each transcript. Colour code: Red indicates significantly upregulated transcripts (FDR ≤ 001 and log_2_ fold change ≥ 1), Blue indicates significantly downregulated transcripts (FDR ≤ 001 and log_2_ fold change ≤ −1), and Grey indicates significantly differential expressed genes, which are not up- or down-regulated (FDR ≤ 001). C). GO-term and pathway enrichment analysis of significantly down-regulated genes in Sb- (left) and Bu-treated cells (right) based on the g:profiler analysis tool. Representative terms are shown. The graphs display q-values and amount of regulated genes for each indicated GO-term and KEGG pathway. See also Figure S6C. D) Cell cycle analysis. Jurkat-E6 cells were treated with Sb or Bu for 48 h. Afterwards, cells were fixed, stained with propidium iodide (PI) and analyzed by flow cytometry. Cell cycle phases were determined using FlowJo software. Histogramms on the left show representative measurements. Donut charts on the right display the amount of cells in the different cell cycle phases as derived from 3 independent experiments. E) Determination of 2-LTR circles. Graph displays the percentage of HIV-1-based 2-LTR circles measured in compound-treated CD4^+^ T-cells. Romidepsin (Ro) was used as internal control. Data are derived from 3 independent experiments and are normalized to vehicle-treated control cells. Error bars display SD. Statistics was performed using the ANOVA-based Tukeys’ multiple comparison test.

### Effect of compound treatment on viral replication steps

More detailed analysis revealed downregulation of HIV entry receptors CD4, CXCR4 and CCR5 as well as changes in T-cell and activation markers CD3, CD25 and CD69, which could be a reason for the observed effects. However, using a virus fusion assay, which allows determination of fused particles with the host cell [41] and thus analysis of the earliest steps of viral replication, we could not determine a significant fusion decline (**Figure EV6D**). Particle fusion in HDACI-treated cells was similar to untreated cells or cells that were treated with the cyclodepsipeptide romidepsin (Ro), which displayed no significant effect on *de novo* infection (**Figure 2C**). Next, we controlled the impact of both inhibitors on T-cell proliferation, as GO-term analysis revealed high regulation of genes involved in the cell cycle. To investigate the impact on cell cycle progression, we treated Jurkat-E6 T-cells with Sb, Bu, or vehicle control and analysed the fraction of cells in G0/G1, G1/S, and G2/M phase by propidium-iodide-based flow cytometry (**Figure 3D**). Defective cell proliferation was already visible during cell culture as reduced medium acidification could be observed for Sb-, and to lower extend, also for Bu-treated cells (decelerated colour change of pH indicator). While treatment of cells with Sb caused a marked block in transition from S to G2/M phase (increase of cells in S-phase of >10 % as well as a decrease of cells in G0/G1 phase of >13 % in comparison to control cells), Bu treatment resulted only in a modest upregulation of both, G1 and G2 phase associated with a decline of cells in S phase (**Figure 3D**). Notably, primary, activated CD4^+^ T-cells showed comparable results; however, effects were not as pronounced as in JurkaT-cells due to significantly lower proliferation rate (data not shown). As a decline in T-cell activation markers as well as an impaired proliferation could indicate a more restrictive, replication suboptimal, intracellular environment, comparable to resting CD4^+^ T-cells, we tested if reverse transcription and associated nuclear import of the preintegration complex (PIC) are hampered. Here, we analyzed RT activity (**Figure EV6E**) and quantified HIV-1 2-LTR circles (**Figure 3E**), which are a surrogate marker for completion of reverse transcription as well as nuclear import of the preintegration complex [42]. Yet, quantitative analysis of RT activity and 2-LTR circles showed no significant Sb- or Bu-related effects.

Thus, we surmized that the main effect of both HDACIs on *de novo* infection might be associated, at least in part, with a perturbed transcriptional regulation. Indeed, correlation of NGS data with the Animal Transcription Factor database (AnimalTFDB) [43] revealed 136 deregulated transcription factors (TFs) and 79 cofactors (CFs) for Sb-treated cells, of which 92 TFs and 37 CFs were significantly upregulated and 44 TFs and 42 CFs were downregulated. For Bu-treated cells, we found 20 deregulated TFs (14 downregulated and 6 upregulated) and 25 CFs (2 up- and 23 down-regulated) (**Figure EV8**). Among the downregulated TFs and CFs were several factors, which have been shown to be either upregulated during HIV-1 infection or to be required for transcription of HIV-1, e.g. ZBED2, FOS, BRCA1 [44–46].

While the changes of transcriptional (co-)factors was less pronounced for Bu-treated cells compared to Sb-treated ones, another observation, which could be associated with transcription of viral genes, drew our attention: We identified two significantly disregulated solute carrier proteins, SLC22A17 and SLC40A1,which play key roles in iron homeostasis and transport (reactome DB [47]; R-HAS-917937; p=0.039). Experimental evidences show that iron is usually actively accumulated in HIV infected cells and seems to be required for HIV-gag expression [48–50]. Earlier studies have shown that the hydroxamic acid bufexamac is capable of chelating iron, which at higher concentrations leads to strong induction of hypoxic conditions in the cell [29]. Thus, bufexamac treatment might impede iron accumulation in the nucleus, leading finally to a reduced p24 level. Indeed, in a preliminary experiment, we could show that the negative effect of bufexamac on *de novo* infection with HIV-1_NL4-3_ could be counteracted by FeCl_3_ in a dose-dependent manner (**Figure EV9**). However, this observation might be based on a variety of cellular processes and therefore requires thorough investigation in future studies. Altogether, further characterization of RNA-Seq data confirms impact of both inhibitors on cell cycle and links decreased de novo infection with impaired transcription of viral genes.

### Global analysis of proteome changes upon HDACI treatment

While analysis of the transcriptome allows determination of ongoing intracellular changes, it usually does not correlate very well with protein abundancies as mRNA levels do not reflect protein half-lives or provide information about posttranslational modifications, which affect protein activity [51,52]. Thus, we additionally performed quantitative, TMT-based proteome analysis of Sb-treated CD4^+^ T-cells 48 h post treatment (**Figure 4A**). In case of bufexamac, we utilized proteomic data from Bu-treated HeLa cells from a former study [29]. For Sb-treated CD4^+^ T-cells, we were able to map 4082 proteins (see also **Supplemental data file**), of which 92 proteins were upregulated (≥ 2 x SD) and 93 proteins were downregulated (≤ 2 x SD) upon treatment, respectively (**Figure 4B**). The bufexamac data set contained 5315 proteins, of which 78 proteins were up- and 127 proteins were down-regulated based on our criteria (≥ 2 x SD and ≤ 2 x SD), respectively (**Figure 4C**). Although, we observed as expected only low correlation on single gene/protein level (only modest correlation between NGS (DEGs) and MS data (R^2^ Sb: 0.2; R^2^ Bu: 0.05)), GO-term and pathway analysis of network associated proteins mirrored the results of the transcriptome analysis, as e.g. terms like “cell cycle” (Sb: Reactome pathways, FDR: 0.0059; Bu: GO-BP, FDR: 0.0338) and “chromatin binding” (Sb, GO-MF, FDR: 0.0138) were significantly enhanced. Among the deregulated proteins, we identified several, which have been previously linked to HIV-1 infection and replication [36] and whose changed abundancies might have an unfavorable effect on viral replication (**Figures 4D, EV10**). For example, proteins like *Rac GTPase activating protein 1* (RACGAP1), *Ubiquitin conjugating enzyme E2C* (UBE2C), *Heterogeneous nuclear ribonucleoprotein U* (HNRNPU), and *3-Hydroxy-3-Methylglutaryl-CoA Synthase 1* (HMGCS1), which have been shown by shRNA/siRNA experiments to be required for HIV-1 replication in Jurkat T-cells [53,54] were downregulated in either Sb- or Bu-treated cells. Also *Dead Box Helicase 5* (DDX5), which has been shown to facilitate HIV-1 replication [55], or the *Histone acetyltransferase P300* (EP300), which acetylates the viral *transactivator of transcription* (Tat) and causes enhanced viral transcription [56] were downregulated in Sb-treated cells (**Figure 4D**). In contrast, proteins like *Ankyrin 1* (ANK1), which acts a an intracellular antiviral agent [57] or *Bromodomain containing 2* (BRD2), which has been identified as an Tat-independent suppressor of HIV-1 transcription [58] were upregulated. Further analysis of deregulated proteins revealed that a large number of these proteins is correlated with funtional association networks as determined by the STRING database [59] (**Figures 4D, EV10**). Overall, proteome datasets corroborate RNA-Seq results and provide further evidence of compound-induced disturbed transcription.

**Figure 4:**
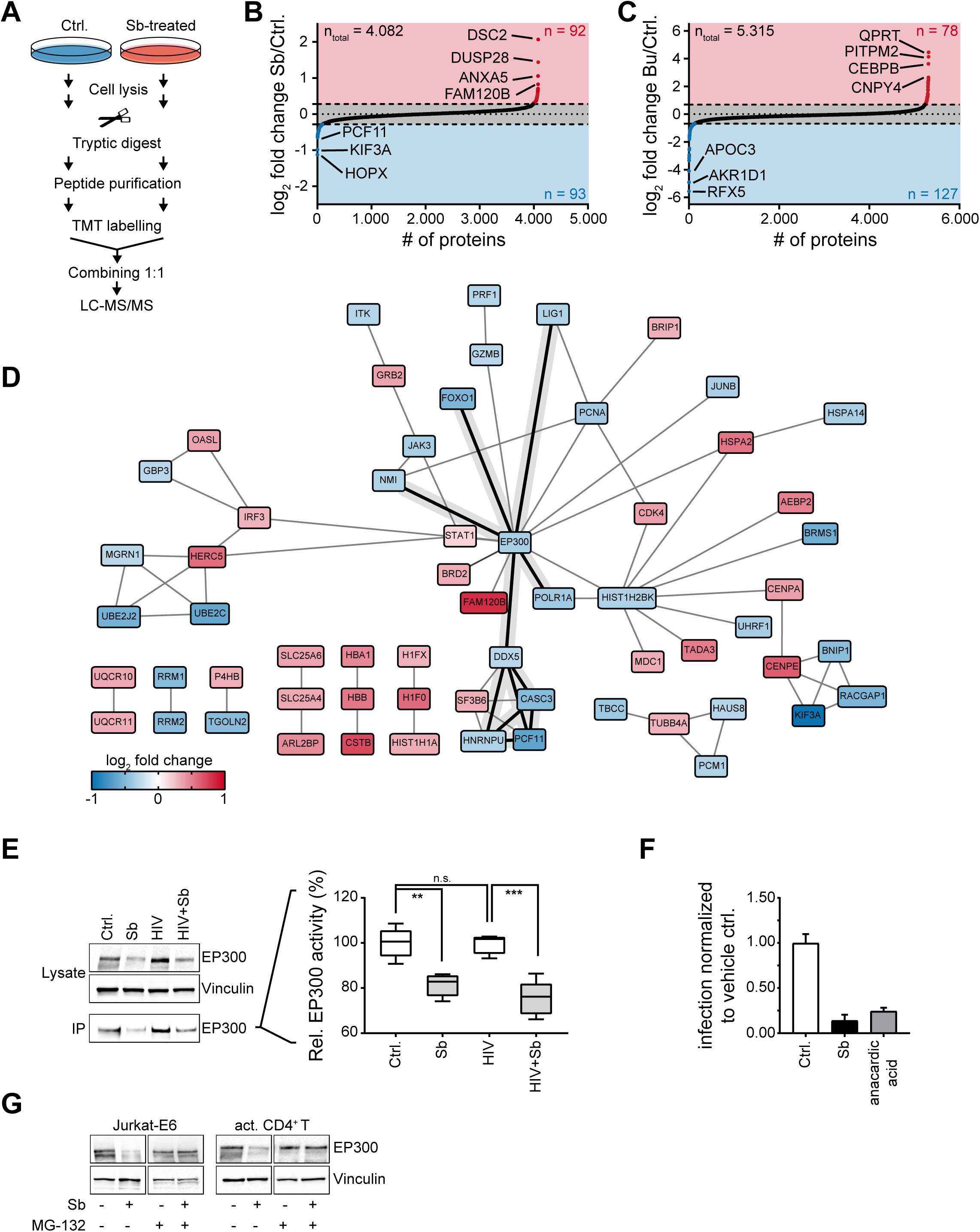
Proteomic analysis of compound-treated cells. A) Schematic illustration of Sb-sample preparation. Cells were challenged for 48 h, lyzed, and subsequently proteins were digested by trypsin. Derived peptides were purified, labeled with isobaric TMT-tags, mixed and analyzed by LC-MS/MS. B) Dot plot displays all identified proteins and the associated log_2_ fold change upon Sb-treament in comparison to control cells. Colored areas represent cut-offs for up-regulation (red; ≥ 2 x SD), down-regulation (blue; ≤ 2 x SD), and no regulation (≤ 2x SD ≥). Numbers (n) of regulated proteins are given on the upper (up-regulated) and lower (down-regulated) right side of the graph. Most changed proteins are indicated by protein name. C) Dot plot displays identified proteins in Bu-treated cells. Data were derived from [29] and were reanalyzed according to similar thresholds as in B. D) Functional interaction network of regulated proteins in Sb-treated cells. Network analysis was performed using STRING database and networks were visualized via cytoscape. The color code displays the measured log_2_ fold changes. Connected proteins within the network, which were down-regulated at protein but not at mRNA level are highlighted (bold black lines). E) Immunoblot of direct cell lysates from treated and HIV-1 infected CD4^+^ T-cells, respectively, as well as of co-immunoprecipitated samples, showing EP300 levels (left). Vinculin served as loading control. Graph on the right shows relative EP300 activity of enriched EP300 (IP samples). Boxes show the lower and upper quartiles, whiskers show the minimum and maximal values, and the line inside the box indicates the median. Statistics was performed using the ANOVA-based Tukeys’ multiple comparison test. F) EP300-associated HIV-1 infection levels. Bars show HIV-1_NL4-3_ infection levels of CD4^+^ T-cells treated either with sodium butyrate or the EP300 inhibitor anarcadic acid (10µM). Data are derived from four experiments and error bars show SD. G) Immunoblots of intracellular EP300 levels in control and Sb-treated cells in presence or absence of proteasome inhibitor MG-132.

### Detailed characterization of Sb-related effects

Comparing the proteome and transcriptome datasets of Sb-treated cells, we noticed that several transcription- and DNA replication-related proteins like EP300, NMI, POLR1A, DDX5, LIG1, FOXO1, CASC3, PCF11, and HNRNPU were profoundly down-regulated on protein level, yet showed no apparent dysregulation on mRNA level (**Figure 4D**). To test whether this observation might be based on protein-ubiquitination and associated proteasomal degradation, we focussed on the acetyl-transferase EP300, which has been shown to be important for enhanced transcription of viral genes [26,27,60]. First, we verified Sb-related EP300 reduction by immunoblotting and analyzed whether a loss of EP300 can be also observed in Sb-treated, HIV-1 infected cells. Activated CD4^+^ T-cells were treated with sodium butyrate or vehicle control only for 24 h and subsequently infected with HIV-1_NL4-3_ and then cultivated for two more days. A profound reduction of cellular EP300 levels was seen upon Sb-treatment, irrespectively of HIV-1 infection (**Figure 4E, left panel**). Notably, the reduction of p24 positive cells upon Sb-treatment was comparable to the treatment of CD4^+^ T-cells with the known EP300 and EP300/CBP-associated factor histone acetyltransferase inhibitor anacardic acid (**Figure 4F**). Additionally, we immunoprecipitated EP300 from cell lysates (**Figure 4E, left panel**) and used the enriched acetyltransferase in combination with the fluorometric SensoLyte HAT (p300) assay (AnaSpec) to probe its activity. Inline with the immunoblot results, significantly reduced EP300 activities were found in precipitates from Sb-treated cells (**Figure 4E, right panel**). Next, we analyzed whether EP300 becomes proteasomally degraded following Sb-treatment. To this end, Jurkat-E6 cells as well as primary, activated CD4^+^ T-cells were challenged for 44 h with sodium butyrate and subsequently cultivated for 4 h in the presence or absence of the proteasome inhibitor MG-132 (**Figure 4F**). Strikingly, immunoblotting of cell lysates showed a recovery of EP300 levels by MG-132 treatment in both cell types, thus corroborating our theory. Very interestingly, recent studies observed a significant lower abundance of butyrate-producing bacteria in the colon/gut of HIV infected individuals [61,62]. It has been surmised that butyrate or more general short-chain fatty acids (SCFAs) may reduce gut inflammation due to induction of regulatory T-cells and modulate activation of antigen-presenting cells [61]. In combination with our observations, one could hypothesize that the effect of butyrate on HIV is thus more direct. Clearly, further research is needed to address the possible linkage between our observations and the rendered microbiome. However, based on our results and the importance of EP300 for viral replication [26,27,56,63–66], it can be assumed that the decline of EP300 together with the very strong effect on cell cycle and the broad modulation of transcription factors highly contributes to the observed reduction of HIV infection in primary, activated CD4^+^ T-cells (**see also Figure EV11**).

### Conclusion

Taken together, our comprehensive screen identifies in particular class I and II HDACs as regulators of HIV-1 gene expression and latency and minimizes the clinical concern of reservoir reseeding by HDACIs. Concurrently, data classify the HDACIs Sb and Bu as novel compounds for efficient reduction of *de novo* HIV-1 infections in primary CD4^+^ T-cells. Furthermore, we provide several evidencies that this reduction could result from compound-mediated rearrangements in the host cell, including deregulation of host cell proliferation and transcriptional control. Additionally, we could show that Sb causes proteasomal degradation of the acetyltransferase EP300. Thus, both compounds might be able to improve existing combined antiretroviral therapy approaches as well as “kick and kill”-strategies – either in combination with other reactivating HDACIs like romidepsin or as single effectors. In particular Sb treatment should be considered in this respect as it not only decreased de *novo infection* of HIV-1, but also allowed reactivation of latent virus and might also support the restoration of the gut microbiome in infected patients.

## Material & Methods

### General reagents and chemical compounds

All standard reagents and chemical compounds were obtained from Carl Roth GmbH + Co. KG/Germany, Sigma Aldrich/Germany, Thermo Fisher Scientific/USA, and TH. Geyer/Germany if not otherwise stated. A list of key resources can be found in **Table 1**.

### Bacterial work

Competent strain Stbl II of *E. coli* was transformed with HIV-1 infectious molecular clone vector pNL4-3 or pCH058/2960 or Blam-Vpr plasmid pMM310 and bacteria were grown in Ampicillin containing TB medium. Subsequently, bacteria were harvested and DNA was isolated with the NucleoBond Xtra Midi Kit (Macherey Nagel) according to manufacturer’s instruction. Concentration of purified DNA was measured with a Nanodrop One^C^ (Thermo Fisher Scientific) and authenticity verified by control digest.

### Standard Cell culture & virus production

Hek293T and Jurkat-E6 cells were cultured in standard medium (DMEM for Hek293T, RPMI for Jurkat-E6) supplemented with 10 % FCS, 100 units/ml penicillin and 0.1 mg/ml streptomycin and cells were cultured at 37°C in 5 % CO_2_. J-Lat cell clones 8.4 and 10.6 were cultured in low endotoxin RPMI (Biochrom GmbH) supplemented with 10 % FCS, 100 units/ml penicillin and 0.1 mg/ml streptomycin and cells were cultured at 37°C in 5 % CO_2_. All cells were confirmed to be mycoplasma negative (Lonza MycoAlert).

For virus production, Hek293T-cells were transfected with pNL4-3 or pCH058/2960 or pNL4-3 and pMM310 (pBlam-Vpr) via polyethylenimine (Polysciences Inc.)[67]. Cells were cultured and 48 h post infection, supernatant was collected, filtered through a 0.45 µm pore size membrane (Merck Millipore/Germany) to remove residual cell debris, and purified by sucrose cushion-based ultracentrifugation (24.000 x g, 4°C, 90 min). Pelleted virus was resolved in PBS (Gibco) and stored at −80°C until further usage. All produced virus stocks were titered by infection of known numbers of relevant targeT-cells under standard experimental conditions, followed by paraformaldehyde (PFA)-based inactivation/fixation of virus infected cells. Subsequently, cells were analyzed by flow cytometry for expression of HIV p24 capsid protein (see below) to identify the fraction of infected cells and to calculate the number of infectious units.

### T-cell isolation and activation

Blood cones (Terumo BCT leukocyte reduction system) containing red blood cells and enriched leukocytes were obtained from the Hospital of the University of Munich, Dept. of Immunohematology, infection screening and blood bank (ATMZH). For isolation of CD4^+^ T-cells, blood cells were diluted with PBS (Gibco) and T-cells were isolated via the Easy-Sep™ Rosette Human CD4^+^ T-cell enrichment kit (Stemcell Technologies, Canada) according to manufacturer’s protocols. Purity of isolated CD4^+^ T-cells was controlled on a FACSverse™ flow cytometer (BD Biosciences) using BD Tritest CD4/CD8/CD3 antibody stain (BD Biosciences). CD4^+^ T-cells were further cultivated and activated for 4 days in RPMI Glutamax medium (Gibco) containing 10 % heat-inactivated FCS (Sigma Aldrich), antibiotics (100 U/ml Penicillin, 100 mg/ml Streptomycin (Merck KgAa)), IL-2 (100 U/ml; Biomol, Germany), and phytohemagglutinin-P (PHA-P) (5 µg/ml; Sigma Aldrich). Subsequently, medium was exchanged to RPMI Glutamax medium (Gibco) containing 10 % heat-inactivated FCS (Sigma Aldrich), antibiotics (100 U/ml penicillin, 100 mg/ml streptomycin (Merck KgAa)), and IL-2 (100 U/ml; Biomol, Germany) without PHA-P and cells were cultivated for another 1-4 days. Again activation was controlled by flow cytometry using anti-human CD4/CD69/CD3 antibody staining solution (BD Biosciences) for identification of activated T-cells.

### Latency reversal assay

J-Lat clones 8.4 and 10.6 were cultivated in presence of increasing concentrations of different HDACI (see also **Table 2**). We tested for each compound a concentration range that spanned the physiological relevant dose or has been used also in other studies [17,29,32,68,69]. The respective highest concentration of solvent (either DMSO or H_2_O) was used as control. 24 h post treatment, J-Lat cells were washed with PBS, fixated and virally inactivated by 1 % PFA for 2 h at RT, and subsequently GFP positive cells were determined by flow cytometry. Datasets were normalized to control.

### Microscopy of HDAC inhibitor treated J-Lat cells

J-Lat cells were freed 24 h post inhibitor treatment from cell debris by low speed centrifugation (23 x g, 5 min, RT) and afterwards harvested by centrifugation (400 x g, 5 min, RT). Cell pellets were re-suspended in culture medium supplemented with 4 % PFA (Electron Microscopy Sciences) and inactivated for 90 min at RT. Subsequently, cells were pelleted (1212 x *g*, 10 min, 4 °C), washed with PBS, and embedded in Prolong™ Diamond Antifade Mountant (Invitrogen) on microscopy glass slides sealed by a glass coverslip. Samples were imaged (brightfield exposure time: 200 ms, GFP exposure time: 100 m) using the Eclipse Ti2 microscope (Nikon) in combination with a DS-Qi2 camera (Nikon) under control of NIS Element AR software (v. 5.0, Nikon). Acquired images with 0.365 µm/pixel resolution were overlayed and analyzed using ImageJ (v. 1.52a) and the ND2 Reader plug-in (Nikon Instruments Inc).

### Cell Viability assay

The viability of primary human CD4^+^ T-cells was assessed after exposure of different concentrations of the respective HDACIs (see also **Table 2**). Concentration range was similar to the latency reversal assay. Activated CD4^+^ T-cells were cultured for 72 h in RPMI medium, supplemented with 10 % heat-inactivated FCS, 100 U/ml penicillin, 100 mg/ml streptomycin and 100 U/ml IL-2 and the respective HDACI. Subsequently, viability of cells was analyzed using the CellTiter-Glo® Luminescent cell Viability Assay (Promega) according to manufacturer’s instructions. Luminescence was measured with a CLARIOStar Plus platereader (BMG Labtech).

### Flow cytometry

Cells were washed with PBS (In case of infection experiments, cells were fixated with PFA prior the PBS wash). Subsequently, cells were either stained with antibody dissolved in PBS containing 3 % albumin (surface markers) or permeabilized with Perm/Wash buffer (BD Biosciences) according to manufacturer’s instruction and stained with the respective antibody (mix) dissolved in Perm/Wash buffer (intracellular targets). Afterwards, cells were washed twice with PBS and analyzed in a FACSVerse and FACSLyric flow cytometer (BD Biosciences), respectively. Data analysis was performed using FlowJo software (TreeStar).

### Verification of increased acetylation

To test global HDACI dependent acetylation, activated T-cells were treated with HDACI as described before and after 72 h total protein acetylation of the cells was determined by flow cytometry using PE-coupled anti-Acetylated Lysine Antibody (Biolegend). Cells were washed with PBS, fixed with 4 % PFA for 90 min at RT, and permeabilized with Perm/Wash buffer (BD Biosciences) according to manufacturer’s instructions. Subsequently, cells were stained with PE-coupled anti-Acetylated Lysine Antibody or PE-coupled Mouse IgG2b κ Isotype control Antibody (isotype control) (both Biolegend) for 30 min at 4°C. Finally, cells were washed two times with Perm/Wash buffer and resuspended in PBS prior to flow cytometry analysis. Furthermore, we analyzed acetylation levels of Histone H3 in HDACI treated cells using the PathScan® Acetylated Histone H3 Sandwich ELISA Kit (Cell Signaling Technology) as described by the manufacturer. Absorbance was determined via a CLARIOStar Plus platereader (BMG Labtech).

### De novo infection of cells

Activated CD4^+^ T-cells or Jurkat-E6 cells were treated with HDACIs or vehicle controls and cultivated for 24 h. Subsequently, cells were infected with identical amounts of HIV-1 (NL4-3, or CH058) and cultivated for 48 h. Cells were harvested, inactivated/fixated by PFA and then analyzed for expression of HIV p24 capsid protein by flow cytometry using FITC coupled KC57 antibody.

### Transcriptome analysis

RNA was isolated from trizol lysates of cells using Direct-zol RNA Miniprep Kit (Zymo Research) following manufacturer’s instructions. Isolated RNA was additionally purified using Agencourt RNAClean XP beads (Beckman Coulter) following manufacturer’s instructions. After quantification (Nanodrop), the RNA was quality controlled using a Bioanalyzer (Agilent Technologies). Good quality RNA (RIN > 7) was used to generate sequencing libraries by using 500 ng total RNA in a Sense mRNA Seq Libarary Prep Kit V2 for Illumina platforms (Lexogen) following manufacture’s instructions. Libraries were quantified and subsequently sequenced on an Illumina HiSeq 1500 (sequencing mode: 100 nt, single-end). The sequencing data was preprocessed on a Galaxy server (hosted by LAFUGA, Gene Center, Munich). After demultiplexing, the data was trimmed according to Lexogen. The output was mapped to the human genome (hg19) using STAR (v. 2.5.2b-0). Abundant reads were further analyzed using HTSeq-count (v. 1.0.0) and a differential gene expression analysis was performed using DESeq (v. 1.0.19) setting the FDR < 0.01. Adjacent data analysis was performed using Perseus (Version 1.6.5.0) [70], R software environment (Version 3.6.2) [71] in combination with the bioconductor package (Version 3.10), g:Profiler (v. e98_eg45_p14_ce5b097) [40], and GraphPad prism software (v. 7.05).

### Fusion assay

Assay was performed as described in [72]. In brief, cells were infected with HIV-1_NL4-3_ harboring β-lactamase (BlaM)-Vpr. 4 h p.i. cells were harvested by centrifugation (500 x g, 5 min) and washed with CO_2_-independent medium supplemented with 10 % FCS (Thermo Fisher Scientific). Afterwards, cells were loaded with CCF2 dye using the LiveBLAzer FRET – B/G Loading Kit (Thermo Fisher Scientific) according to manufacturer’s instruction. After incubation over night at RT, cells were fixed with PFA and analyzed by flow cytometry

### Cell cycle analysis

For cell cycle analysis, Jurkat-E6 cells were treated with HDACIs as described before and cultivated for 24-72 h (only data from 48 h cultivation are shown). Following, cells were washed with PBS and fixed in 70 % ice-cold ethanol for at least 2 h. Fixed cells were pelleted (3500 x g, 5 min), re-suspended in cold PBS containing 0.25 % Triton X-100, and incubated for 15 min on ice. Cells were pelleted again and re-suspended in PBS containing 10 µg/ml RNase A (Sigma Aldrich) and 20 µg/ml propidiumiodide (Sigma Aldrich) and stained for 30 min at RT in the dark. Afterwards, cells were washed with PBS and analyzed in a FACSLyric flow cytometer (BD Biosciences). Data analysis was performed using FlowJo software (TreeStar).

### Analysis of RT activity

CD4^+^T-cells were treated with Sb, Bu, or vehicle control and infected 24 h post treatment with HIV-1_NL4-3_. Cells were cultured for 24 h and intracellular RT activity was determined with colorimetric reverse transcriptase assay (Sigma Aldrich) according to manufacturer’s instruction.

### Determination of 2-LTR circles

2-LTR circles were measured as described [73,74]. Briefly, total DNA from HIV infected cells was prepared using the Blood & Cell Culture DNA Mini Kit (Qiagen). The amount of 2-LTR circles was analyzed by quantitative PCR. DNA was amplified using the PowerUp SYBR Green Master Mix (Thermo Fisher Scientific) and forward 5’-CTAACTAGGGAACCCACTGCT-3’ and reverse 5’-GTAGTTCTGCCAATCAGGGAAG-3’ primers. Amplification was monitored using the Quantstudio 3 Real time cycler (Thermo Fisher Scientific). ß-globin (primers forward: 5’-CCCTTGGACCCAGAGGTTCT-3’; reverse: 5’-CGAGCACTTTCTTGCCATGA-3’) was used as an internal control to normalize total DNA. Data was processed using QuantStudio Design and Analysis software (v. 1.4.3) (Thermo Fisher Scientific).

### MS sample preparation

CD4^+^ T-cells were treated with Sodium butyrate as described before. Vehicle-treated cells were used as control. 48 h post treatment, cells were washed with PBS and directly lysed in modified RIPA buffer (50 mM Tris-HCl (pH 7.5), 150 mM NaCl, 1 mM EDTA, 1 % NP-40 and 0.1 % sodium deoxycholate, supplemented with Complete protease inhibitor mix (Roche). Lysates were mixed with 1/10 volume of 5 M NaCl to release chromatin-bound proteins and incubated for 15 min on ice. Subsequently, lysates were homogenized by sonication (12 × 5 sec with 5 sec pause between cycles, 15 W), and proteins were precipitated overnight at −20 °C by the addition of eight volumes of ice-cold acetone. Acetone precipitates were redissolved in 8 M urea (6 M urea, 2 M thiourea) supplemented with 1 mM DTT for reduction of disulfide bonds and protein concentration was determined by Quick-start Bradford assay (Bio-Rad). Proteins were alkylated with 5.5 mM chloracetamide (45 min, RT) in the dark. The protein solution was eight-fold diluted with Hepes buffer (50mM, pH 7.5) and digested with Pierce Trypsin/LysC Protease Mix (1:100 w/w; Thermo Fisher Scientific) over night. Digestion was terminated by trifluoroacetic acid (TFA) (final concentration 1 %). Resulting peptide mixture was cleared by centrifugation (2,500 x *g*, 5 min) and loaded onto reversed-phase C18 Sep-Pak columns (Waters), pre-equilibrated with 5 ml acetonitrile (ACN) and 2 × 5 ml 0.1 % TFA. Peptides were washed twice with 5 ml 0.1 % TFA and 5 ml H_2_O before elution with 3 ml of 50 % ACN. Purified peptides were dried in a vacuum concentrator (Eppendorf) and re-solubilized in 100 mM triethyl ammonium bicarbonate (TEAB) buffer. Peptide concentration was determined by Bradford assay and 100 µg peptides per condition (control and treated sample) were labeled each with 0.2 mg TMT reagent (TMTduplex Isobaric Mass Tagging Kit; Thermo Fisher Scientific) according to manufacturer’s instructions (see also [75]). Labeling reaction was quenched with 5 % hydroxylamine and equal amounts of labelled peptides were mixed and subsequently dried by vacuum concentration. Peptides were re-dissolved in 0.1 % TFA, concentration was determined by using a Nanodrop One^C^ photospectrometer (Thermo Fisher Scientific), and 100 µg of TMT-labeled peptide mix was fractionated via the Pierce High pH Reversed Phase Peptide Fractionation Kit according to manufacturer’s instructions. Each fraction was dried by vaccum concentration and peptides were re-dissolved in MS buffer (0.5 % acetic acid, 0.1 % TFA) and adjusted to concentration of 0.2 µg/µl prior to MS measurement.

### MS analysis

Peptide fractions were analyzed by online nanoflow LC-MS/MS using a Proxeon easy-nLC 1200 system (Thermo Scientific) connected to a Q Exactive HF-X mass spectrometer (Thermo Scientific). Peptides were loaded onto a chromatography column (length 15 cm, inner diameter 75 µm) packed with C18 reverse-phase material (Reprosil-Pur Basic C18, 1.9 µm, Dr. Maisch GmbH) and eluted by a 106 min gradient (5-40% ACN/H_2_O and 0.1% formic acid). Eluting peptides were ionized by electro spray ionization and injected into the mass spectrometer under the following conditions: 2.0 kV spray voltage, no sheath/aux/sweep gas flow rate, 275°C capillary temperature, and funnel RF level of 40. The Q Exactive HF-X was operated by Xcalibur (v. 4.1.31.9) and Tune (v. 2.9) in data dependent mode. MS1 scans were performed at a resolution of 60,000 (200 m/z), in TopN mode (N=12) and a scan range of 350-1,800 m/z. The AGC target for the scans was 5e5, maximum injection time 50 ms. For MS2 peptides were fragmented at a normalized collision energy of 38 by Higher-energy collisional dissociation (HCD). MS2 scans were executed at a resolution of 45,000 (200 m/z), an AGC target of 1e05 and a maximum injection time of 105ms, isolation window of 0.7 m/z, a fixed first mass of 110 m/z. The minimum AGC target was 4.5e3, dynamic exclusion 20 s and ions with charge states 1 or >7 were excluded from analysis.

MS data were analyzed with MaxQuant (v. 1.6.10.43). All TMT pairs were quantified and MS/MS spectra were searched against the human Uniprot FASTA database (release November 2019) to identify corresponding proteins. The false-discovery rate (FDR) was fixed to a threshold of 1% FDR at peptide and protein level and all peptide identifications were filtered for length and mass error. Cysteine carbamidomethylation was searched as a fixed modification. Statistical analysis was performed using the R software environment (http://www.r-project.org/) in combination with the proteus R package: TMT data [76]. Further data analysis was performed using the STRING database (v. 11.0) [59] and data were visualized using Cytoscape (v. 3.4.0) [77].

### Immunoplots

Cells were treated with compound or vehicle control for 48-72 h. For MG-132 based proteasomal inhibition, cells were additionally co-treated with 1 µM MG-132 (final conc.) 4 h before harvest. Afterwards, cells were lysed in modified RIPA buffer, and cell debris was removed by centrifugation (5 min, 20.000 x g). The cleared lysate was mixed with NuPage 4x LDS sample buffer (Thermo Fisher Scientific) incl. 100 mM DTT. Samples were separated on a NuPage 3-8% Tris-Acetate gel (Thermo Fisher Scientific), blotted onto nitrocellulose membrane and analyzed for EP300 and vinculin (loading control) on a digital blot scanner (Vilber Fusion FX) using p300 (clone NM11, Invitrogen) and vinculin (abcam) primary antibodies and HRP-coupled goat-anti-mouse antibody (Thermo Fisher Scientific).

### EP300 Activity assay

To measure p300 activity in treated cells, cell were lysed with modified RIPA buffer and EP300 was co-immunoprecipitated using magnetic Sure-bead Protein G (Bio-Rad)-coupled p300 (clone 2033, Invitrogen) antibody. After 4 h incubation at 4°C, beads were washed 3x with lysis buffer and purified P300 enzyme was directly used in combination with the SensoLyte HAT (p300)-Assay-Kit (AnaSpec), according to manufacturer’s instruction. Samples were measured on a ClarioStar Plus plate reader (BMG labtech).

## Ethics statement

All work with infectious virus or HIV latent cell lines (J-Lat) was performed in a S3** facility. Work with HIV has been approved by the Bavarian government (**AZ 55.1GT-8791.GT-2-1206-11**). Blood cells were derived exclusively from anonymized healthy donors. Usage of blood cones was approved by the ethics committee of the LMU München, Munich, Germany with the project No.: 17-202-UE

## Data deposition

RNA-Seq data have been deposited in NCBI’s Gene Expression Omnibus [78] with the identifier GSE146854 (https://www.ncbi.nlm.nih.gov/geo/query/acc.cgi?acc=GSE146854).

The MS proteomics data have been deposited to the ProteomeXchange Consortium (http://proteomecentral.proteomexchange.org) via the PRIDE partner repository [79] with the data set identifier PXD017909.

## Acknowledgements

J-Lat clones 8.4 and 10.6, were obtained through the NIH AIDS Reagent Program from Eric Verdin[28]. HIV-1 infectious molecular clones pNL43 and pCH058.c/2960 as well as the Blam-Vpr plasmid pMM310 were obtained through the NIH AIDS Reagent Program from Dr. Malcolm Martin [80] and Dr. John Kappes & Dr. Christina Ochsenbauer [35] and Dr. Michael Miller [81]. We would like to thank the Dept. of Transfusion Medicine of the University Hospital Munich for providing blood cones. We are also grateful to Dr. Alexander Graf from the Gene Center Munich for help with GEO. This work was supported by grants from the Else Kröner-Fresenius-Stiftung (2016-A134) to C.S. and by LMUexcellent funding of the LMU Munich to C.S. and H.B.

## Author contributions

Contribution: L.C., A.Z. and C.S. performed experiments and analyzed data. T.L. and C.C. carried out MS analysis. J.PM. and H.B. performed RNA-Seq and helped with data analysis. O.T.K. provided essential equipment and gave advice for virological experiments. C.S. designed the study. C.S., L.C. and A.Z. wrote the manuscript. All authors reviewed the manuscript

## Conflict of interest

The authors declare that they have no conflict of interest.

## Expanded View Figures

**Expanded View Figure 1:**
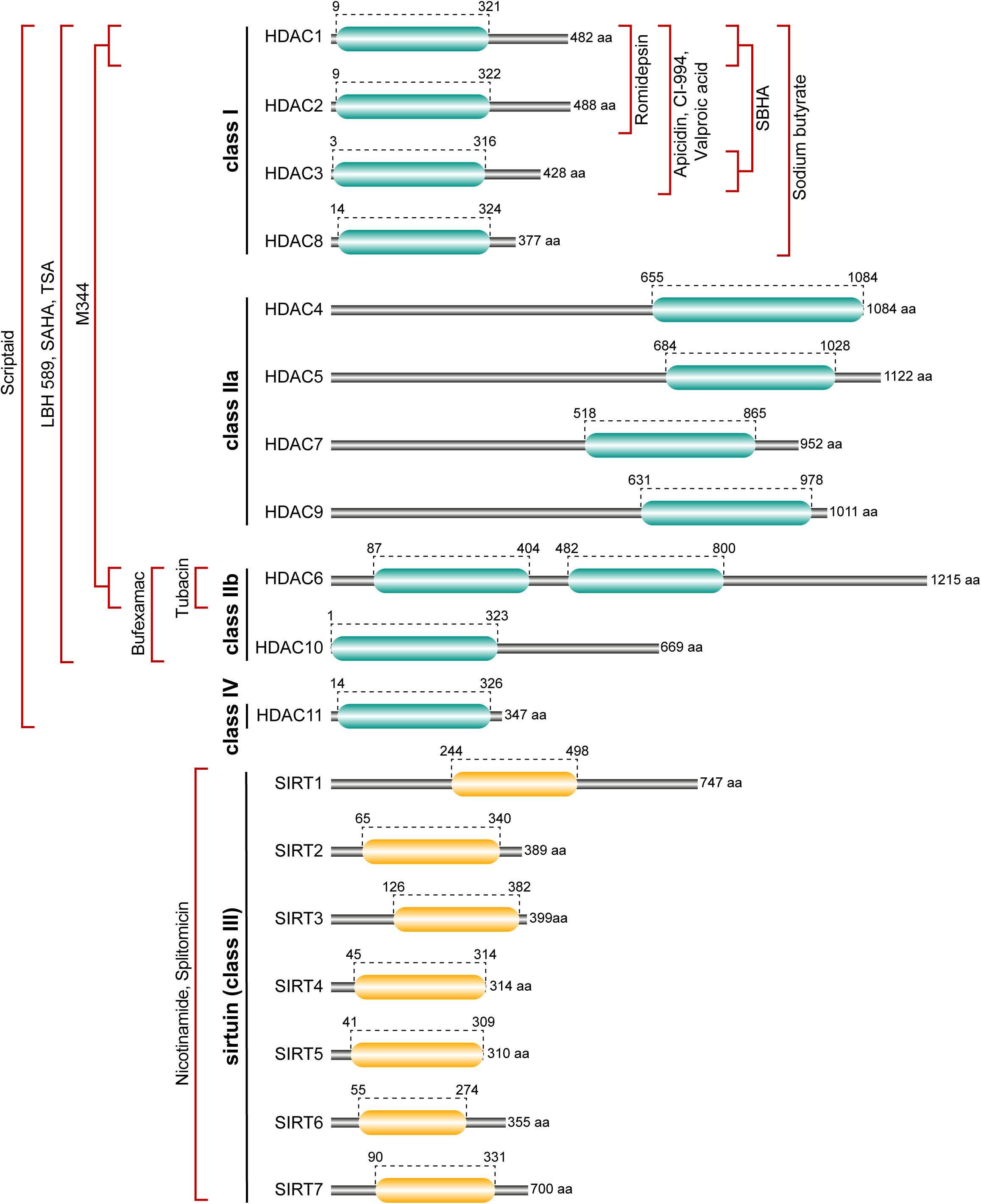
Schematic overview of HDACIs used in this study and effective range. Active domain(s) of human deacetylases are highlighted as follows: green box = deacetylase domain; yellow box = sirtuin type deacetylation domain. Amino acid numbers indicate start and end of the domain(s) as well as of the whole protein. Data based on the UniProt database (www.uniprot.org).

**Expanded View Figure 2:**
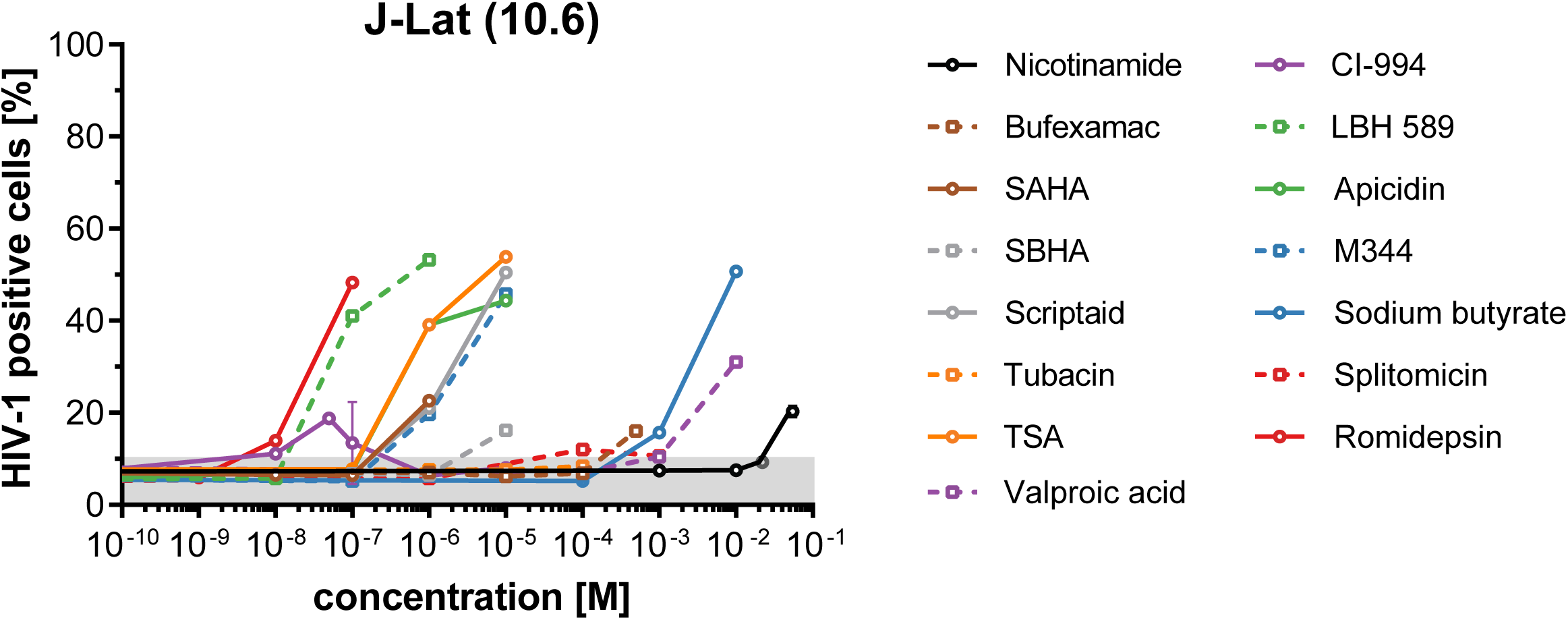
HIV-1 latency reversal in J-Lat cells (clone 10.6). Cells were treated with increasing inhibitor concentrations for 24 h. Subsequently, percentage of reactivation was determined by GFP-based flow cytometry. Data are derived from three independent experiments. Error bars show SD. See also Figure 1A.

**Expanded View Figure 3:**
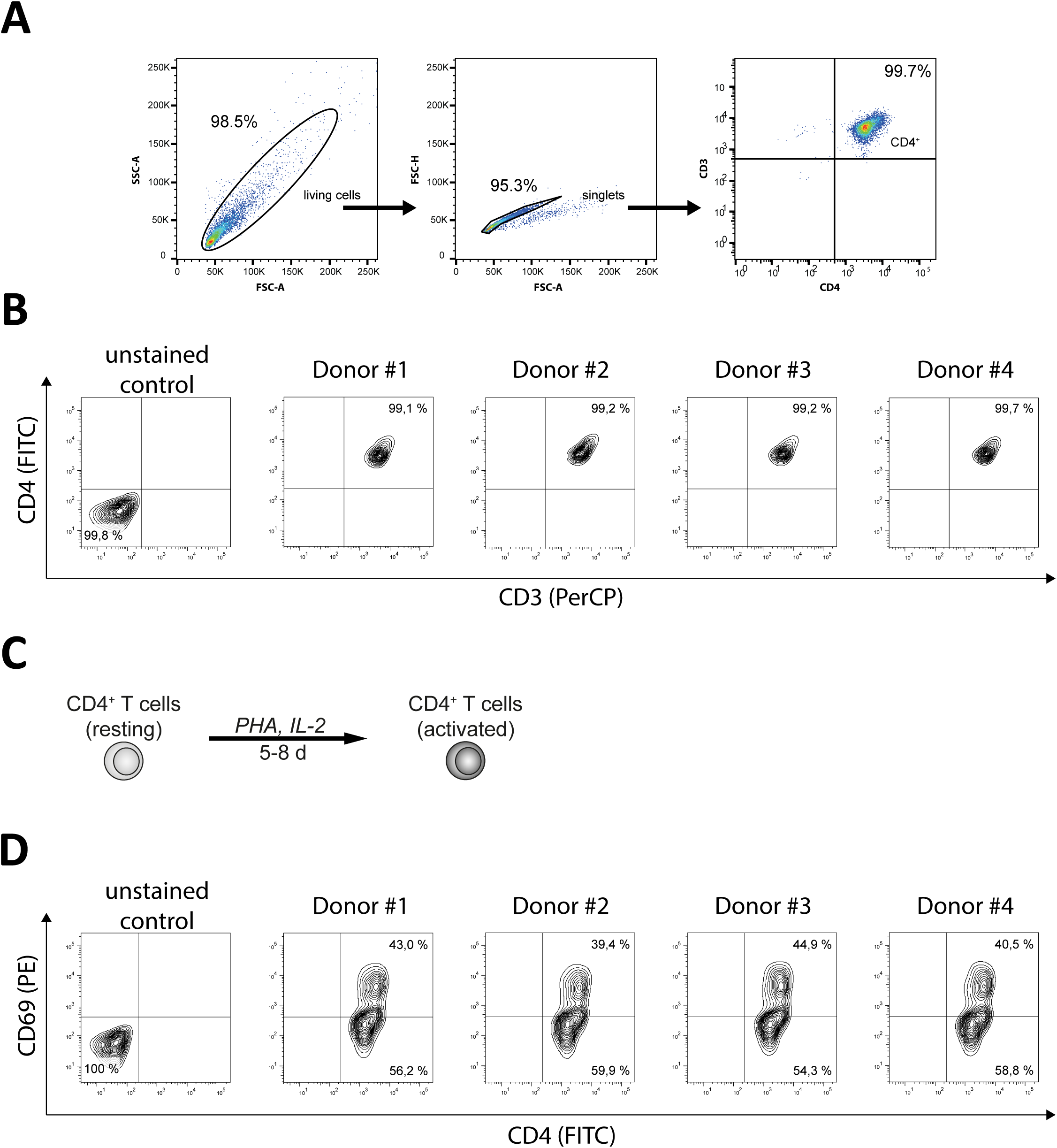
Isolation and activation of CD4^+^ T-cells. A) CD4^+^ T-cells were isolated from blood of healthy donors using RosetteSep™ technology. Subsequently, purity of isolated cells was analyzed by fluorescence-based flow cytometry using anti-CD3 / -CD4 staining. FACS plots display the gating strategy for living cells (lymphocytes; left), singlets (middle), and enriched CD4^+^ T-cells (right). Numbers display the percentage of cells within the respective gate. B) Representative FACS plots of CD4^+^ T-cell isolation based on the gating strategy in A from four different donors, which were used later in four-donor-pools. C) Schematic illustration of T-cell activation. Isolated CD4^+^ T-cells (resting) were cultivated in the presence of PHA and IL-2 for 5-8 days to gain activated CD4^+^ T-cells. D) Representative FACS plots of the isolated cells from B after activation. CD69 was used as activation marker.

**Expanded View Figure 4:**
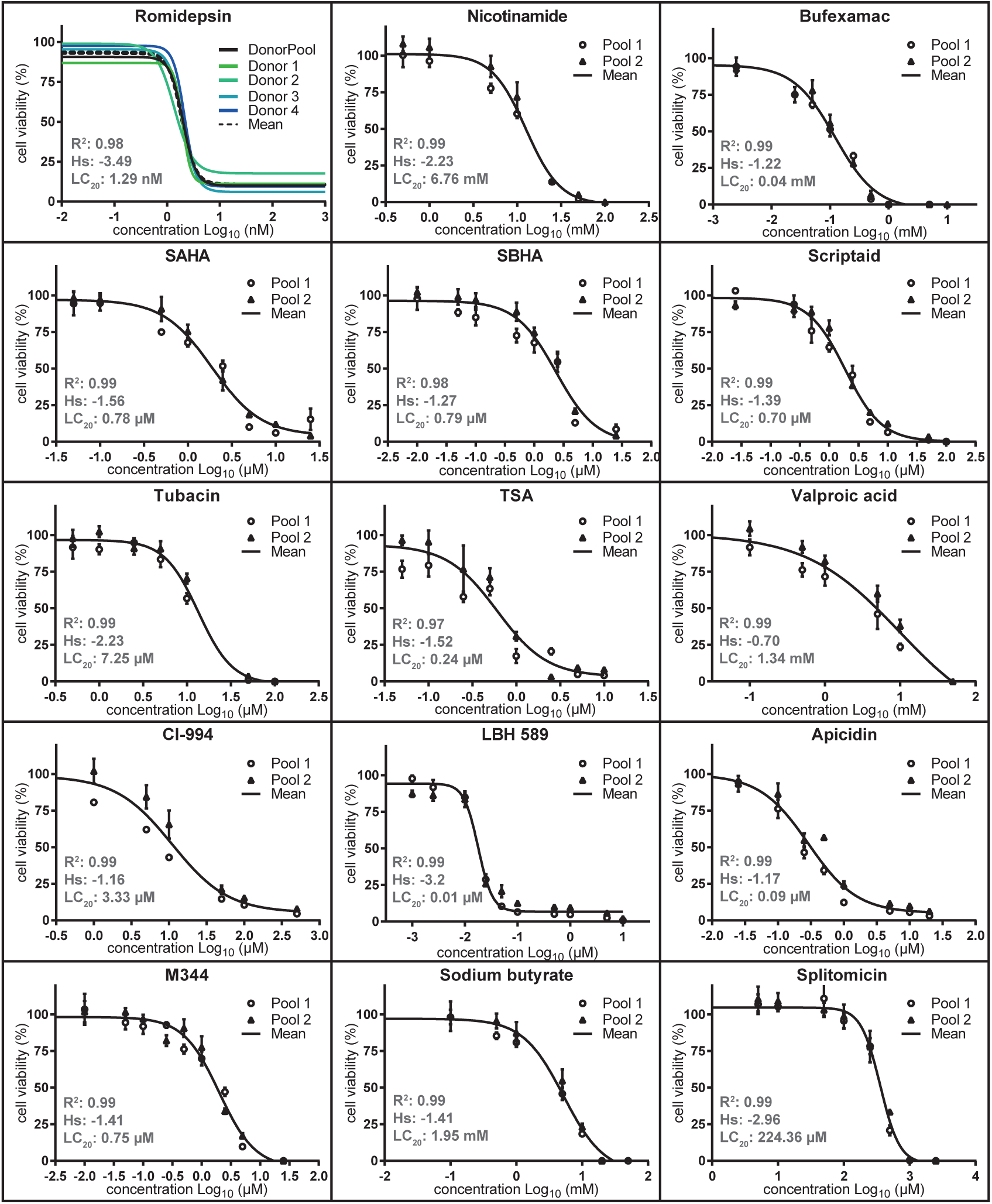
Toxicity / cell viability screen. Activated CD4^+^ T-cells were cultivated in the presence of different concentrations of the respective HDACIs for 72h. Afterwards, viability of cells was determined using a luminescence-based viability assay. Besides individual data points the fitting curve of all measurements, R^2^ value, Hill slope (Hs), and lethal concentration of 20% (LC_20_) is displayed. Data are derived from three independent experiments each with 2 independent four-donor pools. Error bars show standard deviation (SD). For Romidepsin the mean of three independent experiments as well as concentration-dependent viability curves of 4 individual donors are shown (upper left graph).

**Expanded View Figure 5:**
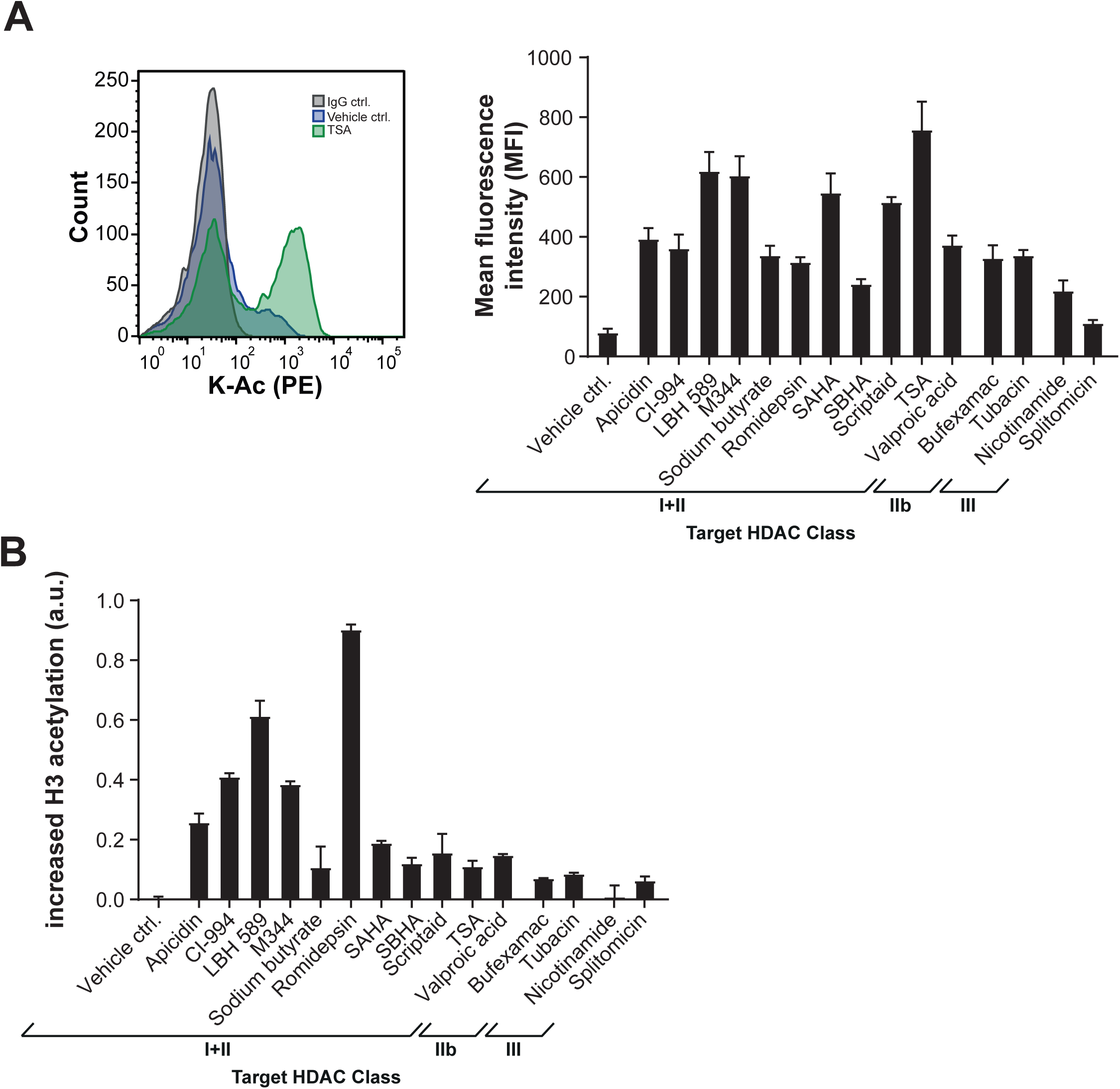
Verification of compound activity. A) Left: Exemplary FACS plot showing global protein acetylation levels of vehicle-treated (blue) or TSA-treated (green) T-cells. Specificity of anti-acetyl antibody was tested using TSA-treated cells in combination with IgG-control antibody (grey). Right: Bar diagram shows the mean fluorescence intensities (MFI) of cellular acetylation levels after HDACI treatment. B) Bar diagram displays the increased histone H3 acetylation as determined by ELISA assay and normalized to vehicle-treated control cells. Data in A and B are derived from three independent experiments. Error bars show standard deviation (SD).

**Expanded View Figure 6:**
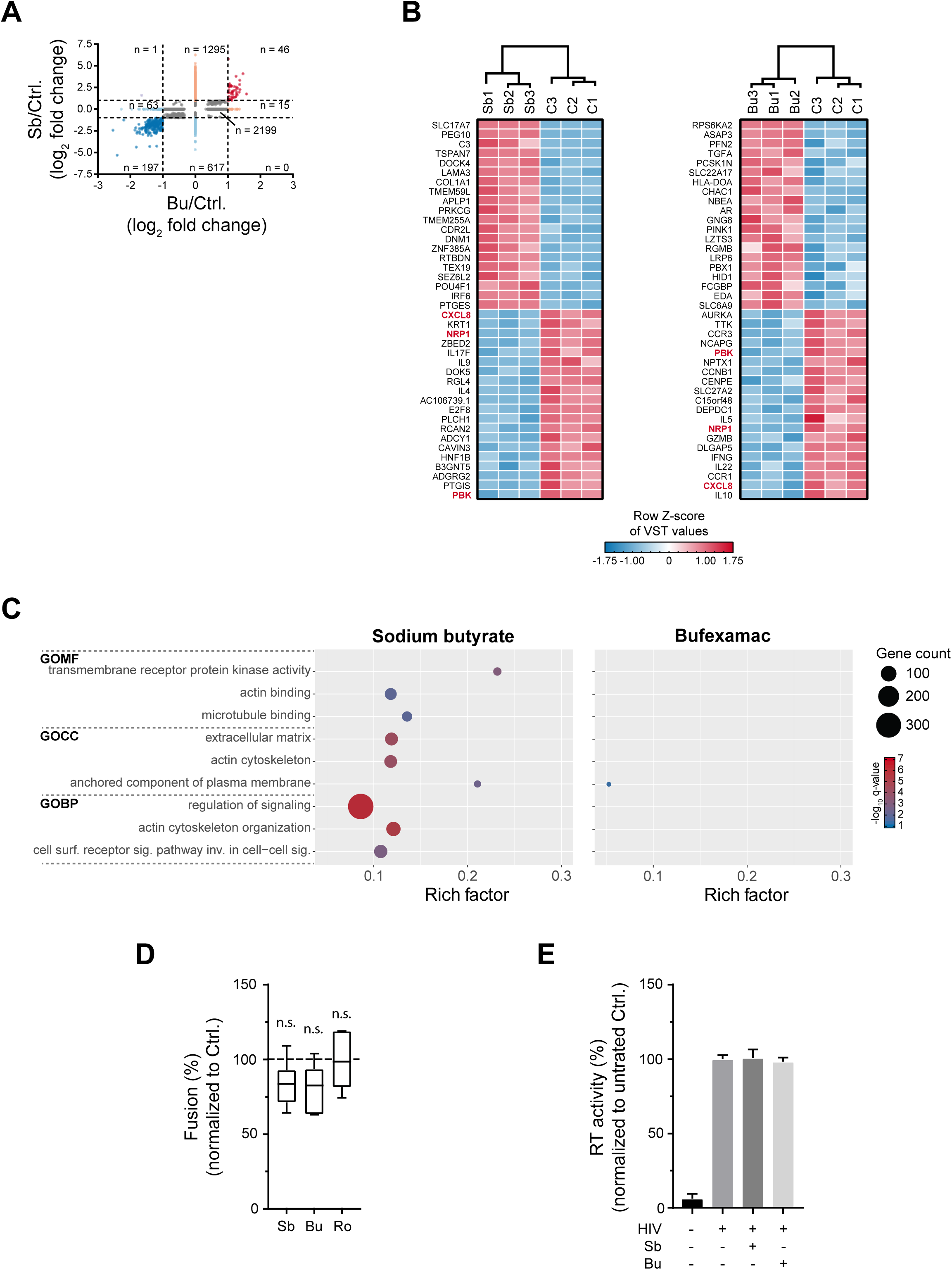
Further analysis of transcriptome data. A) Correlation plot of RNA-Seq data from Sb- and Bu-treated cells. Only transcripts are shown, which could be assigned to an ENSEMBL ID. DEGs with log_2_ fold change > −1 and < 1 are depicted in grey. Regulated DEGs (log_2_ fold change ≥ 1 or ≤ −1) are indicated as follows: upregulated in only one condition (orange), upregulated in both conditions (red), downregulated in only one condition (light blue), downregulated in both conditions (dark blue), and contrary regulated (magenta). n = number of DEGs belonging to the respective group. B) Heatmaps showing the 20 most up- or down-regulated genes in Sb-treated (left) and Bu-treated (right) cells. Colour intensity shows row z-score of variance stabilized transformation (VST) values. Top 20 genes that have been identified in both treatment conditions are indicated with red letters. C) GO-term-analysis of up-regulated genes. Left: Sb-treatment, Right: Bu-treatment. Representative terms are shown. The graphs display q-value and amount of regulated genes for each indicated GO-term. See also Figure 3C. D) Viral particle fusion assay. Box-Whisker plot displays the percentage of fusion of viral particles with the target cell. Datasets were normalized to untreated control cells and were derived from three independent experiments. Boxes show the lower and upper quartiles, whiskers show the minimum and maximal values, and the line inside the box indicates the median. Statistics was performed using one-way ANOVA-based Dunnetts’ multiple comparison test. Sb: Sodium Butyrate, Bu: Bufexamac, Ro: Romidepsin. E) RT activity assay. Graph displays RT activity in dependency of HDACI treatment as analyzed in HIV-1 infected CD4^+^ T-cells. Data were derived from four independent experiments; error bars show SD.

**Expanded View Figure 7:**
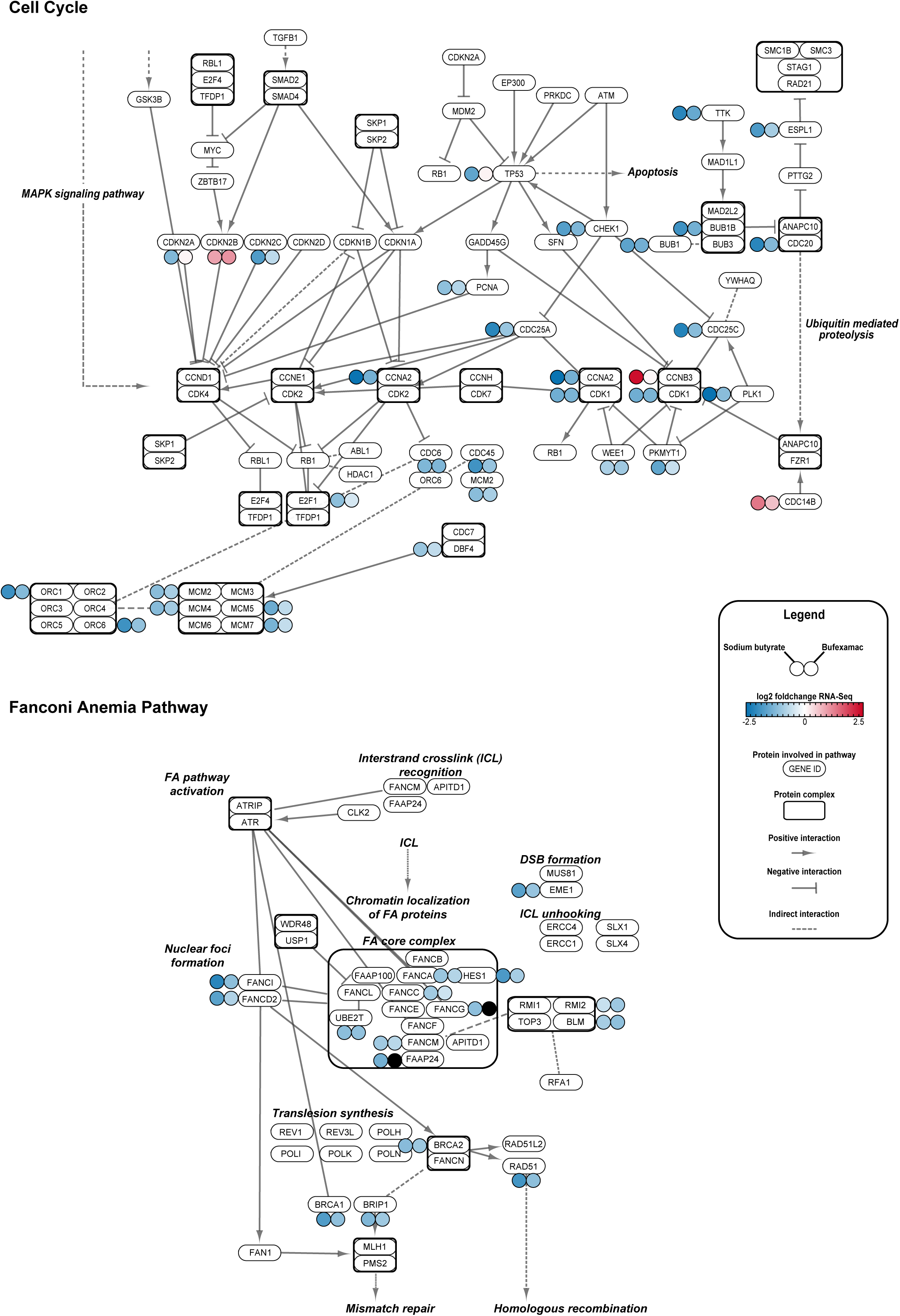
KEGG analysis of significantly up- and down-regulated genes. Schematic representation of the KEGG-terms “Cell cycle” (upper diagram) and “Fanconia anemia pathway” (lower diagram). Genes/Proteins involved in the cellular pathway are displayed in white color. Protein complexes are additionally surrounded by a black frame. Positive interactions (e.g. induction of expression, activation) are shown by arrows, negative interactions (e.g. repression) are displayed by bar-headed arrows. Indirect or unknown interactions are indicated by dashed lines. The log_2_ fold change of significantly DEGs is shown in the immediate vicinity of the respective gene symbol in small circles (left side: abundance of genes in Sb-treated samples in comparison to control and right: abundance of genes in Bu-treated samples in comparison to control. Black circles: Gene has not been identified in the related sample).

**Expanded View Figure 8:**
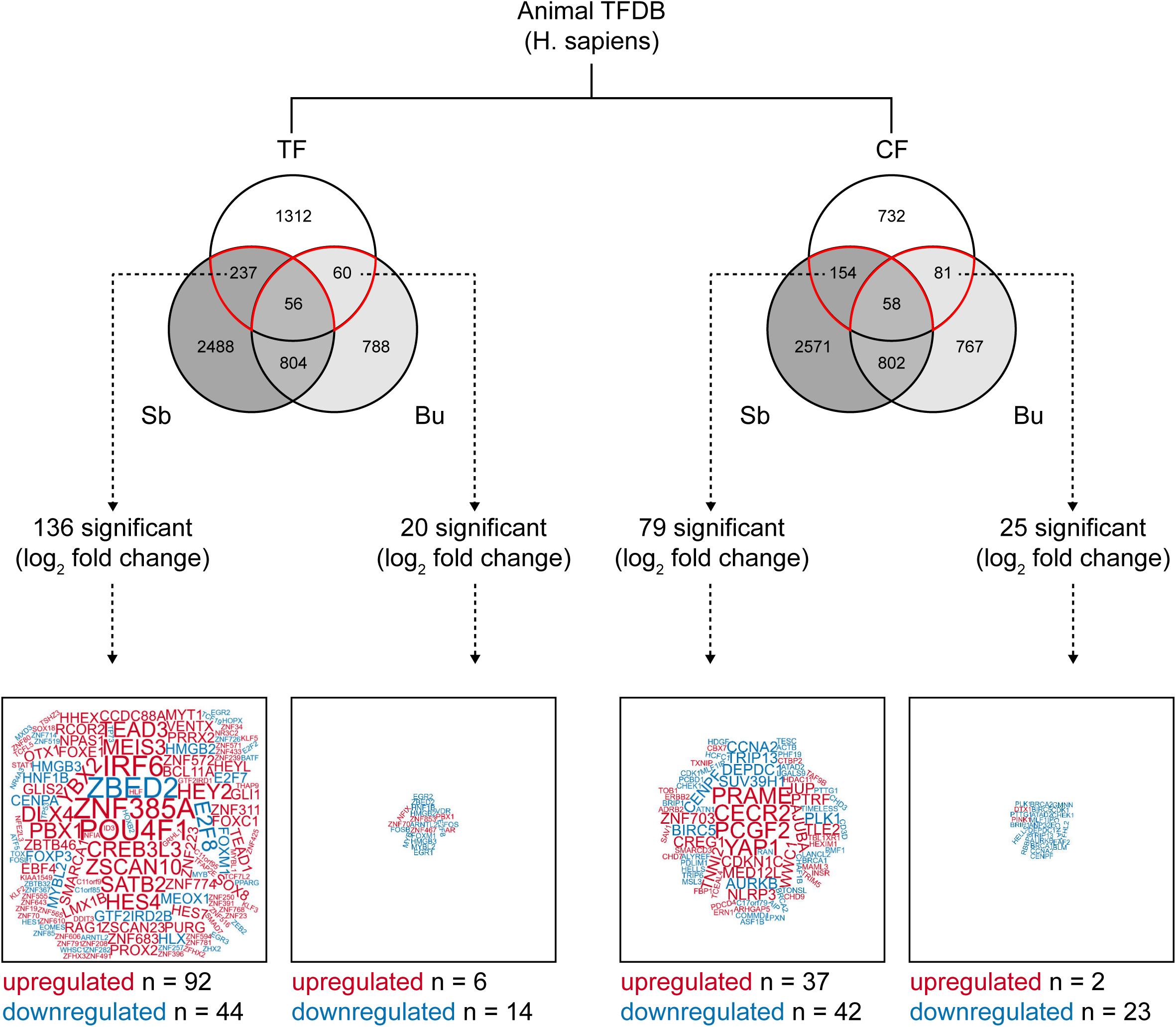
Deregulated transcriptional factors and cofactors. Venn diagrams (upper part) show the number of transcription factors (TF; left side) and transcriptional cofactors (CF; right side) listed in the Animal TFDB [43] and the overlap with DEGs from Sb- (dark grey) and Bu-treated (light grey) cells. TFs and CFs, which were significantly deregulated (based on log_2_ fold change) are represented in word clouds (lower part). Up-regulated genes are colored red, down-regulated genes are colored blue. Letter size reflects log2 fold change. Word clouds are generated with the free online tool *http://wortwolken.com*.

**Expanded View Figure 9:**
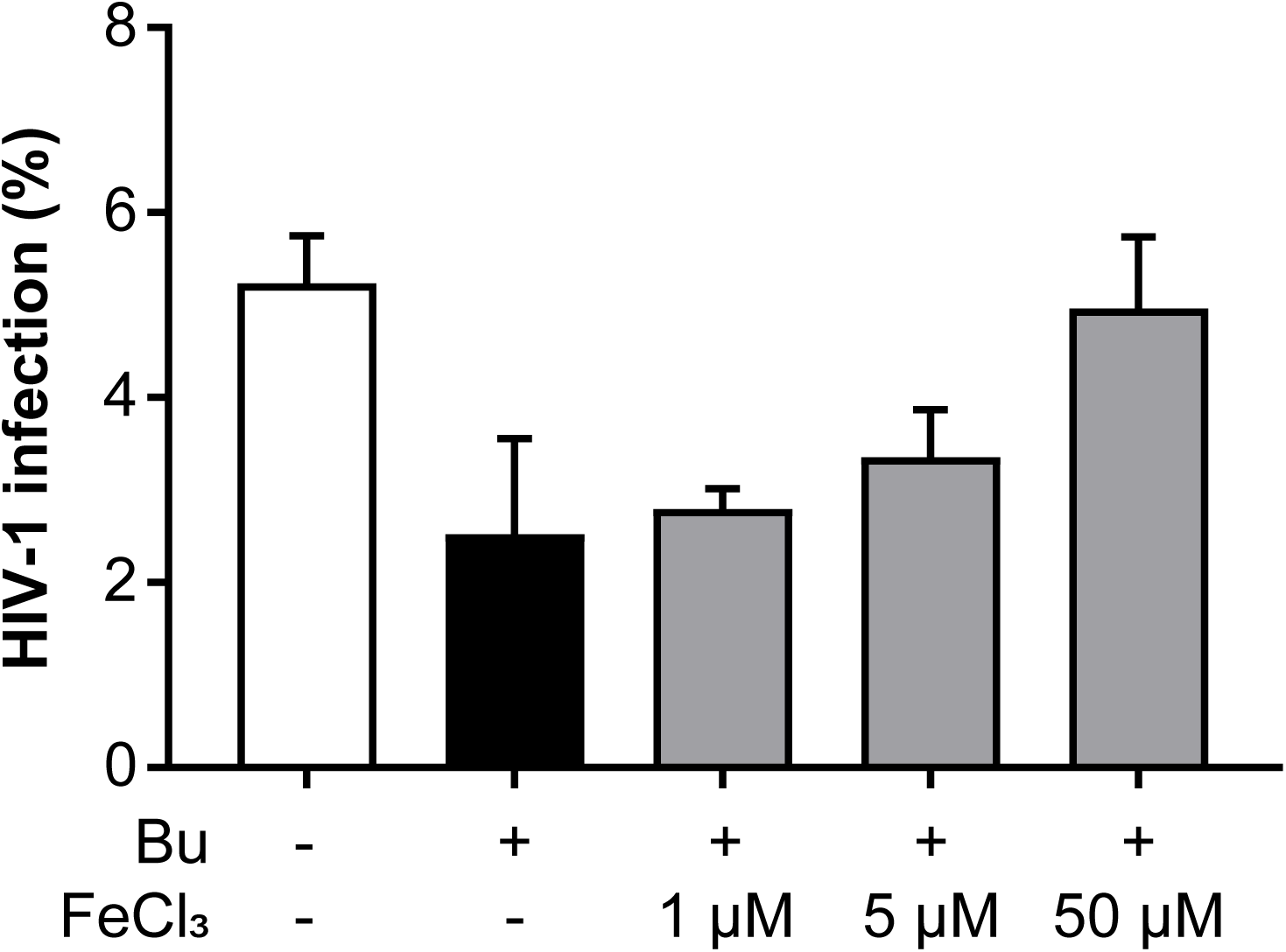
Reversal of bufexamac-induced reduction of p24 expression by iron. CD4^+^ T-cells were treated with bufexamac in presence of increasing concentrations of iron (FeCl_3_). 24 h post treatment, cells were infected with HIV-1_NL4-3_ and further cultured for another 48 h. Subsequently, cells were analyzed for p24 via flow cytometry. Untreated cells were used for control.

**Expanded View Figure 10:**
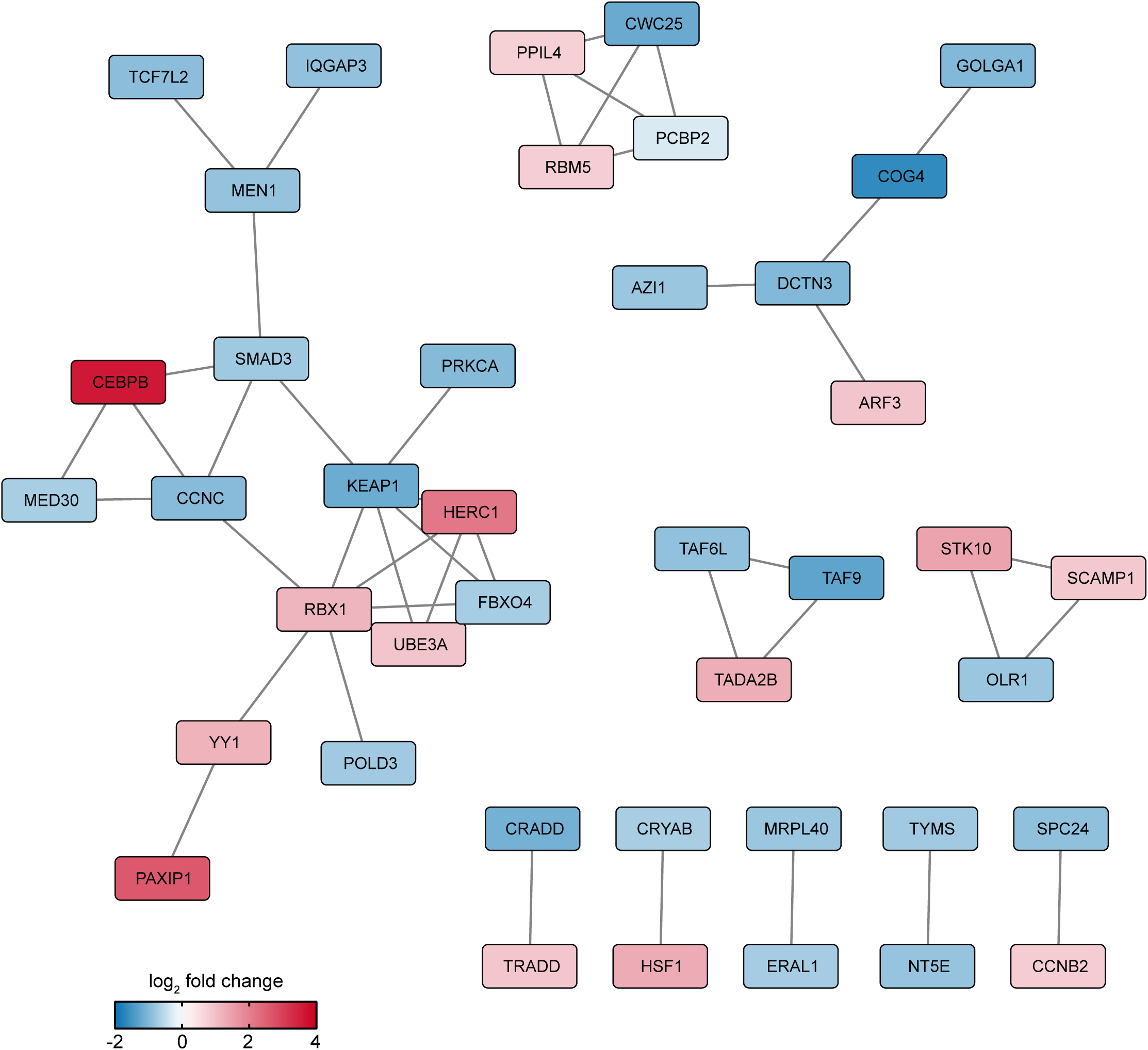
Functional interaction networks of Bu-regulated proteins. Network analysis was performed using STRING database and networks were visualized via cytoscape. The color code displays the log_2_ fold changes as derived from the bufexamac dataset [29].

**Expanded View Figure 11:**
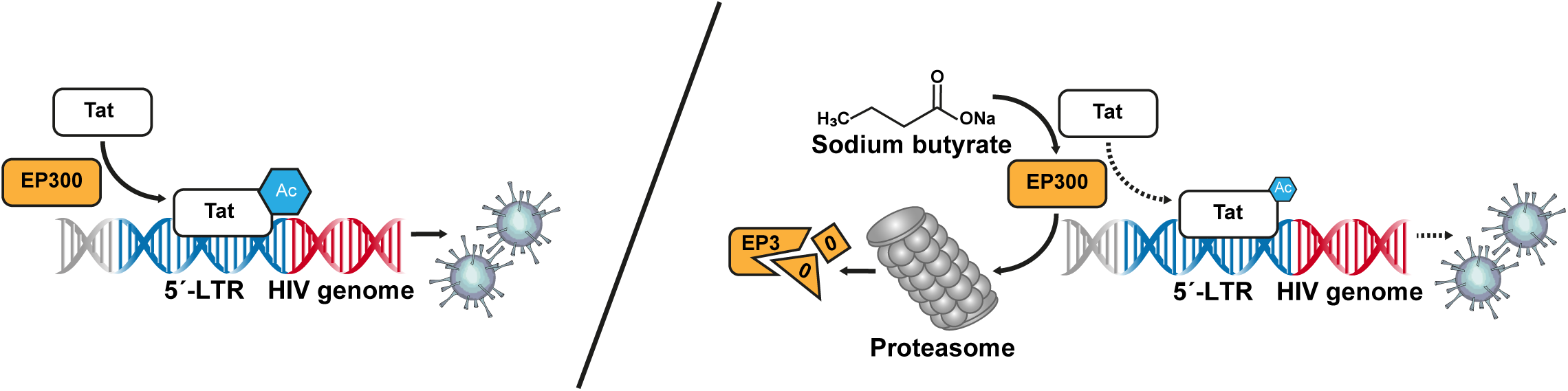
Data-based model of the molecular events underlying the Sb-treatment. At standard infection conditions, the acetyltransferase EP300 is recruited by the viral TAT protein and enhances its transcriptional activity by acetylation (left). In presence of sodium butyrate, EP300 becomes proteasomally degraded, causing a reduced acetylation of TAT itself and at the same time a reduced transcription of viral proteins (right).

